# Structural conservation of Lassa virus glycoproteins and recognition by neutralizing antibodies

**DOI:** 10.1101/2022.09.26.509601

**Authors:** Hailee R. Perrett, Philip J. M. Brouwer, Jonathan Hurtado, Maddy L. Newby, Judith A. Burger, Lin Liu, Joey H. Bouhuijs, Grace Gibson, Terrence Messmer, John S. Schieffelin, Aleksandar Antanasijevic, Geert-Jan Boons, Max Crispin, Rogier W. Sanders, Bryan Briney, Andrew B. Ward

**Affiliations:** Department of Integrative, Structural and Computational Biology, The Scripps Research Institute, La Jolla, CA 92037, USA; Department of Immunology and Microbiology, Scripps Research, La Jolla, CA 92037, USA; Center for Viral Systems Biology, Scripps Research, La Jolla, CA 92037, USA; School of Biological Sciences, University of Southampton, Southampton, SO17 1BJ, United Kingdom; Department of Medical Microbiology and Infection Prevention, Amsterdam University Medical Centers. Location AMC, University of Amsterdam, Amsterdam Infection & Immunity Institute, Amsterdam, 1105 AZ, The Netherlands; Complex Carbohydrate Research Center, University of Georgia, Athens, GA 30602, USA; Department of Pediatrics, Tulane University School of Medicine, New Orleans, LA 70112, USA; Department of Chemical Biology and Drug Discovery, Utrecht University, Utrecht, 3584 CG, The Netherlands; Department of Microbiology and Immunology, Weill Medical College of Cornell University, New York, NY 10021, USA

**Keywords:** Lassa mammarenavirus, Lassa fever, arenavirus, structure-based vaccine design, cryoEM, neutralizing antibody, prefusion glycoprotein

## Abstract

**Summary:** Lassa fever is an acute hemorrhagic fever caused by the zoonotic Lassa virus (LASV). The LASV glycoprotein complex (GPC) mediates viral entry and is the sole target for neutralizing antibodies. Immunogen design is complicated by the metastable nature of recombinant GPCs and the antigenic differences amongst LASV lineages. Despite the sequence diversity of GPC, structures of most lineages are lacking. We present the development and characterization of prefusion-stabilized, trimeric GPCs of LASV lineages II, V, and VI, revealing structural conservation despite sequence diversity. High-resolution structures and biophysical characterization of GPC in complex with GP1-A antibodies reveal their neutralization mechanisms. Finally, we present the isolation and characterization of a novel trimer-preferring neutralizing antibody belonging to the GPC-B competition group with an epitope that spans adjacent protomers and includes the fusion peptide. Our work provides molecular detail information on LASV antigenic diversity and will guide efforts to design pan-LASV vaccines.

**Highlights:**

- Structural characterization of soluble glycoproteins from four Lassa virus lineages.
- MAb 12.1F, belonging to the GP1-A cluster, inhibits matriglycan and LAMP-1 binding.
- GP1-A mAbs show glycan-dependence with 19.7E demonstrating lineage-dependent binding.
- A novel trimer-preferring NAb S370.7 targets the GPC-B epitope.

## Introduction

The ongoing SARS-CoV-2 pandemic emphasizes the importance of pandemic preparedness for zoonotic pathogens, which—through climate and anthropogenic variables that increase the landscape suitability for zoonotic transmission—cause approximately 75% of infectious disease in humans (Carlson et al., 2022; Gebreyes et al., 2014). Since its identification in 1969, the Old World arenavirus Lassa (LASV) has caused endemic Lassa fever disease in West Africa. While most cases appear to be asymptomatic (McCormick et al., 1987), an acute hemorrhagic fever can develop leading to high case-fatality ratios often exceeding 25% among patients showing clinical symptoms (Akpede et al., 2019; Ilori et al., 2019; Monath, 2019). LASV is most often transmitted to humans from spillover events with its near-ubiquitous reservoir host *Mastomys natalensis*, which is otherwise known as the natal multimammate rat. Transmission more rarely occurs via nosocomial infection (Dan-Nwafor et al., 2019) and sexual transmission post-recovery (Thielebein et al., 2022). Because of its substantial genomic variability, LASV is subdivided into seven lineages (I-VII; Ruo et al., 1991; Whitmer et al., 2018; Yadouleton et al., 2020). This variability increases the difficulty of developing robust diagnostics, likely resulting in an underrepresentation of LASV’s disease toll (Bowen et al., 2000; Kafetzopoulou et al., 2019; Siddle et al., 2018). There are no efficacious treatments or vaccines for this disease except the controversial off-label use of ribavirin and supportive care (McCormick et al., 1986). Owing to this, the World Health Organization and the Coalition for Epidemic Preparedness Innovations recognize the need for increased LASV research and development efforts given its pandemic potential (Mehand et al., 2018) and have supported early-stage vaccine development and corresponding clinical trials (Gouglas et al., 2019).

The glycoprotein complex (GPC) is the sole viral protein on the surface of LASV and presents the target for neutralizing antibodies (NAbs; Robinson et al., 2016; Watanabe et al., 2018). GPC, which is expressed as a single polypeptide and proteolytically processed by site-1 protease (Rojek et al., 2008), is a trimer of heterodimers and is comprised of the receptor-engaging subunit GP1 and transmembrane-spanning subunit GP2. Approximately 25% of the GPC molecular weight is attributable to the highly conserved 11 or 12—lineage-depending—potential N-linked glycosylation sites (PNGS) per monomer (Eichler et al., 2006; Watanabe et al., 2018). As a result, GPC carries a dense glycan shield which contributes to LASV’s evasion of neutralizing humoral immune responses (Sommerstein et al., 2015). Similar to HIV-1 Env, LASV GPC features a cluster of oligomannose-type glycans (Watanabe et al., 2018) that function as attachment factors and enable LASV’s infection of immune cells via DC-SIGN (Goncalves et al., 2013). Shedding of GP1 during acute disease in humans has been observed and is thought to act as an immune decoy given the conformational variability between GP1 presented as part of GPC and soluble, shed GP1 (Branco and Garry, 2009; Branco et al., 2010; Hastie et al., 2017). LASV exploits two host cell receptors to infect human cells. Host cell attachment is mediated by matriglycan moieties on α-dystroglycan which interact with residues on the GPC trimer apex (Acciani et al., 2017; Katz et al., 2022; Sheikh et al., 2022; Willard et al., 2018). Upon macropinocytosis and trafficking of LASV through the endosomal compartments (Oppliger et al., 2016), GPC undergoes a pH-dependent switch allowing binding to endosomal receptor lysosomal-associated membrane protein 1 (LAMP-1). Putative residues for LAMP-1 binding involve the histidine triad and supporting GP1 residues (Cohen-Dvashi et al., 2015; Israeli et al., 2017).

The largest anti-LASV antibody isolation study to date, which yielded 113 cloned human monoclonal antibodies (mAbs) from memory B cells of LASV survivors, defines the canonical competition groups: GP1-A, GPC-A, GPC-B, and GPC-C (Robinson et al., 2016). X-ray crystallography studies with GPC-B mAbs revealed their epitopes bridged two protomers at the base of the GPC trimer, making contacts with the N-terminal loop, T-loop, HR1 and HR2 helices, and the fusion peptide (Hastie et al., 2017, 2019). Cryo-electron microscopy (cryoEM) structures have shown the GPC-A mAbs target an epitope that extends between the GP1 and GP2 subunits between the N79, N89, N99, N224, and N365 glycans and bind in a three Fab per GPC trimer occupancy (Enriquez et al., 2022). Both antibody competition groups have been shown to lower the fusogenicity of the GPC and limit binding to LAMP-1. Molecular details on the GP1-A competition group, which includes mAbs 12.1F, 19.7E, and 10.4B, as well as the GPC-C group, which consists only of mAb 8.9F, are currently lacking.

Previous structural work of a ligand-free, native-like GPC has been made difficult by the instability of the trimeric ectodomain (Hastie et al., 2017) and inefficient cleavage when introducing stabilization mechanisms (Gorman et al., 2022; Schlie et al., 2010; Willard et al., 2018; Zhu et al., 2021). Published structural information of the GPC in its prefusion conformation is mostly limited to GPC from the lineage IV Josiah strain in complex with antibodies (Enriquez et al., 2022; Hastie et al., 2017, 2019; Katz et al., 2022), although a recent study describes GPC from lineage I (Buck et al.). Our recent work demonstrates fusing GPC to the I53-50A component of the computationally designed I53-50 nanoparticle (Bale et al., 2016) stabilized the trimeric conformation of GPC (Brouwer et al., 2022). In line with the generation of I53-50 nanoparticles presenting glycoproteins of HIV-1, SARS-CoV-2, and RSV, GPC-I53-50 nanoparticles assembled efficiently upon mixing of GPC-I53-50A and the pentameric subunit I53-50B (Brouwer et al., 2019, 2021; Marcandalli et al., 2019). Display of GPC on I53-50 nanoparticles has demonstrated success in eliciting NAb responses *in vivo*; yet the full nanoparticle system complicates structural analysis.

Here, we utilize the I53-50A subunit as a scaffold to generate and characterize GPC trimers of LASV genotypes beyond the prototypical lineage IV Josiah. We focus on lineages of public health concern including lineage II one of the most common lineages which circulates widely in southern Nigeria; lineage V, which circulates in Mali and has decreased pathogenicity compared to lineage IV; and lineage VI, a newly described lineage isolated from a nosocomial infection in Togo with comparable pathogenicity to lineage IV (Mateo et al., 2022; Safronetz et al., 2013). Establishing a single particle cryo-electron microscopy (cryoEM) GPC pipeline allowed us to generate unliganded high-resolution structures of these GPC trimers, revealing structural commonalities and subtle differences between these geographically distinct lineages. In addition, we present the structures of GPC in complex with NAbs 12.1F and 19.7E, adding molecular definition to the mechanism of neutralization of these GP1-A mAbs and their different neutralization phenotypes. Finally, we describe the isolation and structural characterization of a novel trimer-preferring mAb from a Sierra Leonean Lassa fever survivor, providing additional molecular information for the GPC-B epitope cluster. This work not only expands our structural knowledge of the different GPC lineages and their NAb epitopes but also enables investigation of lineage antigenicity at the molecular level—key steps towards the development of a pan-LASV vaccine.

## Results

### Engineering stable prefusion LASV GPC trimers of different lineages

As LASV has known antigenic differences and limited humoral cross-reactivity (Ruo et al., 1991; Whitmer et al., 2018; Yadouleton et al., 2020), we first assessed the sequence conservation of LASV’s GPC across 300+ known sequences. While the GPCs have highly conserved sequences in receptor binding sites and PNGSs, there is notable variability (Fig 1A and B). To study these antigenic distinctions at a molecular level, we expanded our repertoire of recombinant trimeric GPCs. Our previous work demonstrates the ectodomain of GPC (residues 1-424) from the Josiah strain (lineage IV; LIV) can be stabilized as a trimer by fusion to the two-component self-assembling I53-50 nanoparticle (Bale et al., 2016; Brouwer et al., 2022). Further, this immunogen elicited neutralizing humoral responses in rabbits, suggesting it presents epitopes relevant for neutralization (Brouwer et al., 2022). Building off this work, we explore if I53-50A can be used as a trimerization domain to stabilize additional GPCs of diverse lineages. To ensure stabilization of the prefusion state, we introduced the GPCysR4 mutations (Hastie et al., 2017). These mutations comprise the introduction of a disulfide-bond between GP1 and GP2, a proline in the HR1 helix and the replacement of the native site-1 protease cleavage site (Rojek et al., 2008) with a furin cleavage site. The resulting soluble constructs (hereafter referred to as GPC-I53-50As) feature sequences of circulating lineages II (LII; strain NIG08-A41), V (LV; strain Soromba-R), and VI (LVI; strain Togo/2016/7082). GPC-I53-50As were expressed using codon-optimized plasmids in HEK 293F cells and purified as trimer with comparable thermostability (Fig. 1C and D; Fig. S1A and B). Importantly, the trimeric GPC could now be purified in the absence of stabilizing antibodies (Hastie et al., 2017) while showing a high degree of stability and cleavage between GP1 and GP2. Negative stain electron microscopy analysis of purified GPCs demonstrated all constructs formed homogeneous prefusion trimers with an observable smaller density representing the I53-50A trimerization domain (Fig. 1E).

**Fig 1:**
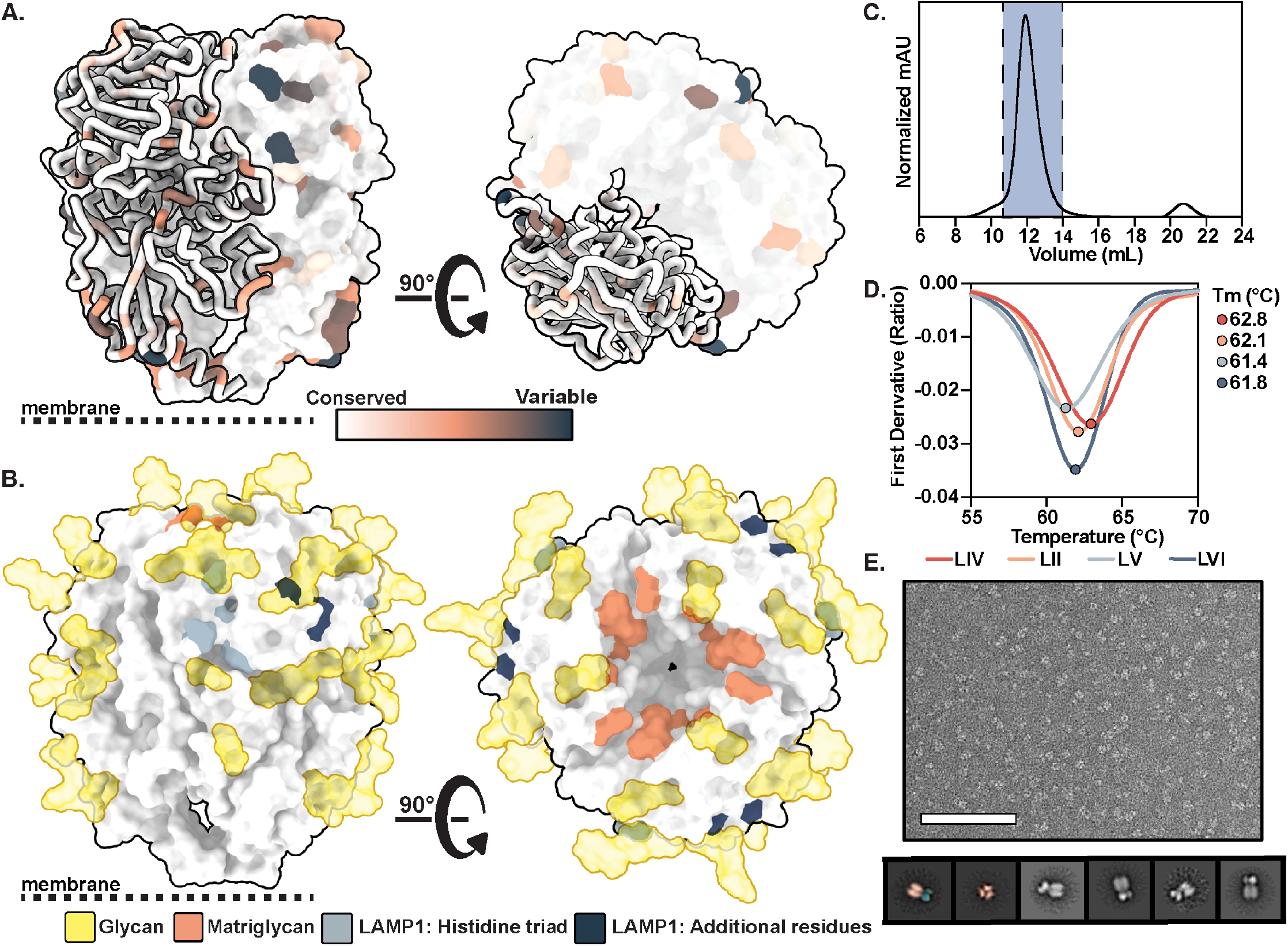
Biophysical characterization of LASV GPC derived from diverse lineages and scaffolded on I53-50A. (A) LASV GPC sequence conservation mapped on ribbon and surface representation of the LIV GPC (PDB 8EJD). Residues with increasing sequence variability are depicted in orange and dark blue, respectively. (B) Glycans from experimental density (gold; EMD-28178) and residues involved in matriglycan binding (orange; Katz et al., 2022) and suspected in LAMP-1 binding (histidine triad in blue, additional residues in gray; Cohen-Dvashi et al., 2015; Israeli et al., 2017) mapped on the surface representation of LIV GPC. (C) Representative size exclusion chromatograph of GPC-I53-50A. The fractions containing GPC-I53-50A trimer are shown in blue. (D) Thermostability of GPC-I53-50As assessed by the inflection point of the curve provided by the ratio of the signal at 350 and 330 nm, as measure by nanoDSF. Circles mark the midpoint of thermal denaturation or melting temperature (*T*_*m*_) of each protein and the values are listed on the right of the graph. Each melting curve is a representative of triplicate curves with melting temperatures within ±0.1°C.

### GPCs from diverse LASV lineages have similar glycan shields

LASV GPC has a highly dense glycan shield (Watanabe et al., 2018) which preferentially envelops GP1 over GP2, resulting in just under half of its surface area being solvent inaccessible (Re and Mizuguchi, 2021). The prototypical LIV GPC has 11 PNGSs on its GPC ectodomain (Eichler et al., 2006; Watanabe et al., 2018), which are thought to contribute to host evasion. LII, LV, and LVI each have an additional PNGS at residue N271 or N272, though this site is uniformly unoccupied (Fig. 2A). Glycan analysis via liquid chromatography-mass spectrometry (LC-MS) revealed a large range of glycan processing states with notable abundance of oligomannose-type glycans near the N- and C-terminal regions of GPC. Complex-type glycans were presented at a higher rate on centrally located PNGSs. Glycan microheterogeneity is pronounced at sites N98/99, N166/167, and N223/224, with each site presenting a mix of oligomannose-, hybrid-, and complex-type glycoforms. This microheterogeneity is largely conserved between lineages (Fig. 2A). The N118/119 site displays near-exclusively complex-type glycans, all of which are fucosylated (Fig. S2A). GPC’s glycan shield features an unusual mannose patch similar to HIV-1 Env (Behrens and Crispin, 2017; Go et al., 2008; Robinson et al., 1987), which is likely caused by steric constraints from neighboring glycan moieties. This restricts access of these PNGS sites to glycan processing enzymes in the endoplasmic reticulum and Golgi apparatus (Watanabe et al., 2018). Previous analysis in a virus-like particle system denotes the mannose patch of LASV GPC as PNGSs N79, N89, N99, N365, and N373 (Watanabe et al., 2018); yet, the lineages presented in Fig. 2A show a large proportion of complex-type glycans presented at N89 and N99. This distinction may be attributable to the expression systems used to generate the VLPs (Madin-Darby canine kidney II cells) or recombinant GPC-I53-50As (HEK 293F cells). With strong conservation of N-linked glycan biosynthesis among mammals, an alternative cause may lie in the oligomerization and cleavage efficiencies of GPC depending on their production as recombinant proteins, presentation on VLPs, or behavior in a native virion context.

**Fig 2:**
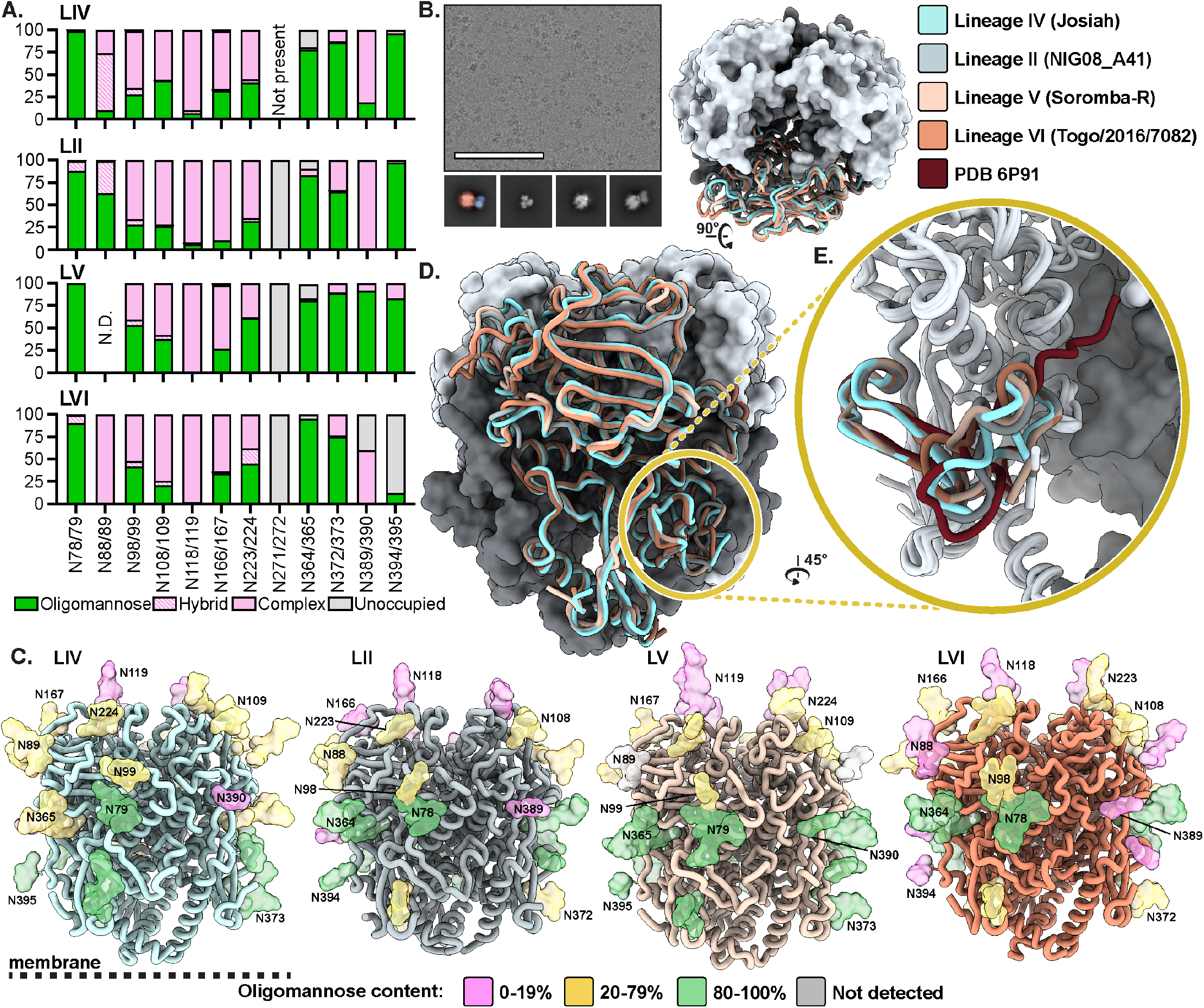
Site-specific glycosylation and structural analysis of LASV GPC from different lineages. (A) Relative quantification of distinct glycan types of GPC determined by LC-MS describe the relative glycan processing state at a particular PNGS. Oligomannose-type glycans are shown in green, hybrid in dashed pink, and complex glycans in pink. unoccupied sites are shown in gray. (B) Representative cryo-electron micrograph of ligand-free GPC-I53-50A. Sample 2D classes are shown below with the leftmost class pseudocolored to indicate the GPC (orange) and I53-50A trimerization scaffold (blue). Scale bar represents 100 nm. (C) Refined atomic models of ligand-free LASV GPC structures of LIV (strain Josiah), LII (strain NIG08-A41), LV (strain Soromba-R), and LVI (strain Togo/2016/7082). Glycans are shown as colored surfaces according to their oligomannose content. Though it is primarily unoccupied on LVI GPC, N394 is colored according to the glycan identity it has when occupied since the glycan was observed in EM data. Access codes are as follows: LIV, PDB: 8EJD, EMD-28178; LII, PDB: 8EJE, EMD-28179; LV, PDB: 8EJF, EMD-28180; and LVI, PDB: 8EJG, EMD-28181. (D) Comparison of models in (C). (E). Comparison of the fusion peptides (LIV and LV residues 260-299; LII and LVII es 259-298) of models in (C) with PDB 6P91 (Hastie et al., 2019), which features the LIV GPC in complex with 18.5C Fab.

### GPCs from diverse lineages demonstrate similar structural features with a distinct fusion peptide conformation

We next assessed whether the LASV lineages present GPCs with distinct structural features. Using single-particle cryoEM, we optimized the conditions for freezing LIV, LII, LV, and LVI GPC-I53-50A trimers (Fig. 2B). Because the GPC is highly glycosylated and has less accentuated features compared to other viral fusion glycoproteins, we found it was difficult to (1) overcome the strong orientation bias of the GPC particle in vitreous ice and (2) align the GPC during data processing when we masked out densities outside of the GPC. Orientation bias likely caused by the interactions of the apex glycans with the air-water interface was relieved by adding a fluorinated detergent to the sample prior to freezing (Fig. S3A). To alleviate poor alignment, we began processing the data using the I53-50A scaffold as a fiducial marker, which facilitated better orientation of the GPC. Combined, these approaches enabled us to resolve the structures of GPC in a reproducible manner and yielded structures of LIV, LII, LV, and LVI GPC trimers at resolutions of 3.8, 3.7, 3.7, and 3.1 Å, respectively (Fig. 1C; Table S1; Fig. S3, Fig. S4).

The GPC-I53-50A constructs recapitulate the known GPC structural features and domain organization (Hastie et al., 2017). GP1 feature the N-terminal β-strands, exterior β-sheet surface, and the interior helix-loop domain. The GP2 subunit demonstrates the canonical HR1a-d helices, T-loop, and HR2 helix. Our GPC-I53-50A shows high similarity, as measured by the root-mean-square deviation (RMSD) of a GPC protomer, to those previously described (0.79 Å with PDB 5VK2 and 0.91 Å with PDB 7PVD; Hastie et al., 2017; Katz et al., 2022). Similarly, when comparing the GPCs of diverse lineages, we observe high homogeneity. Using the prototypical LIV GPC as reference, we note RMSDs of 0.84 (LII), 0.89 (LV), and 0.75 Å (LVI).

The main differences among the GPC lineages were found in flexible loops, most notably the loop (residues 166-181) extending from the β7 sheet prior to the β3 helix. This observed heterogeneity is derived from areas in the EM density of poorer local resolution (Fig. S5), insinuating greater flexibility of the residues in these regions. Consequently, these differences likely do not represent physiologically important conformational epitopes for LII, LV, and LVI GPC.

When comparing our structures to antibody-bound structures reported previously (Hastie et al., 2019), we observe a substantial difference in fusion peptide conformation (Fig. 2E). In the ligand-free structures of GPC-I53-50As, the fusion peptide appears to flexibly occupy the space enclosed by the HR1a helix of the same protomer and the HR1d and HR2 loop of its adjacent protomer. In contrast, previously described crystal structures of GPC bound to 18.5C, 37.7H, and 25.6A of the GPC-B competition group (Hastie et al., 2017, 2019) and 25.10C of the GPC-A competition group (Enriquez et al., 2022) show the fusion peptide occupies the same approximate area, yet extends inwards and reaches toward the apex of the trimer near the GP1 C-terminal domain (Fig. 1E, S2B). This conformational difference increases the buried surface area of residues 260-276 of the fusion peptide from 598 to 621 Å^2^ upon 18.5C binding, for example, and lowers the solvent accessibility of the fusion peptide. Both the antibody-bound and unbound structures show the fusion peptides adopting a near identical conformation starting at the fusion loop (residues 277-299).

### GP1-A epitope mAbs 12.1F and 19.7E neutralize by blocking receptor binding

While the GPC-A and GPC-B antibody interactions with GPC have been studied in detail (Enriquez et al., 2022; Hastie et al., 2017, 2019), molecular details of the GP1-A antibodies have so far remained elusive. Although 12.1F and 19.7E are both members of the described GP1-A competition cluster (Robinson et al., 2016), these mAbs have distinct genetic features. Whereas the heavy chain (HC) and light chain (LC) of 12.1F are derived from the IGHV4-34*01 and IGKV3-11*01, respectively, the germline HC and LC of 19.7E are IGHV3-74*02 and IGKV1-5*01. The VH genes of 12.1F HC and LC are 8.8 and 7.6% somatically hypermutated, respectively, based on the sequences publicly available (Patent WIPO: WO2018106712A1).

To identify differences between 12.1F and 19.7E at the phenotypic level, we analyzed the GPC binding and neutralization of these mAbs to a broad panel of LASV lineages (Fig. 3A and B). Using our suite of stable GPC-I53-50As, we performed biolayer interferometry (BLI) experiments and observed marked differences between the binding behavior of 12.1F and 19.7E among the lineages (Fig. 3A; Fig. S6). When comparing the on-rate of IgG binding to immobilized GPC, we observed the LIV GPC-I53-50A had the highest overall binding efficiency to the tested NAbs. This finding makes sense as the LIV GPC was used as the capture antigen during mAb isolation and both patients from whom the B cells were derived were from Sierra Leone where LIV LASV dominates (Manning et al., 2015; Robinson et al., 2016).

**Fig 3:**
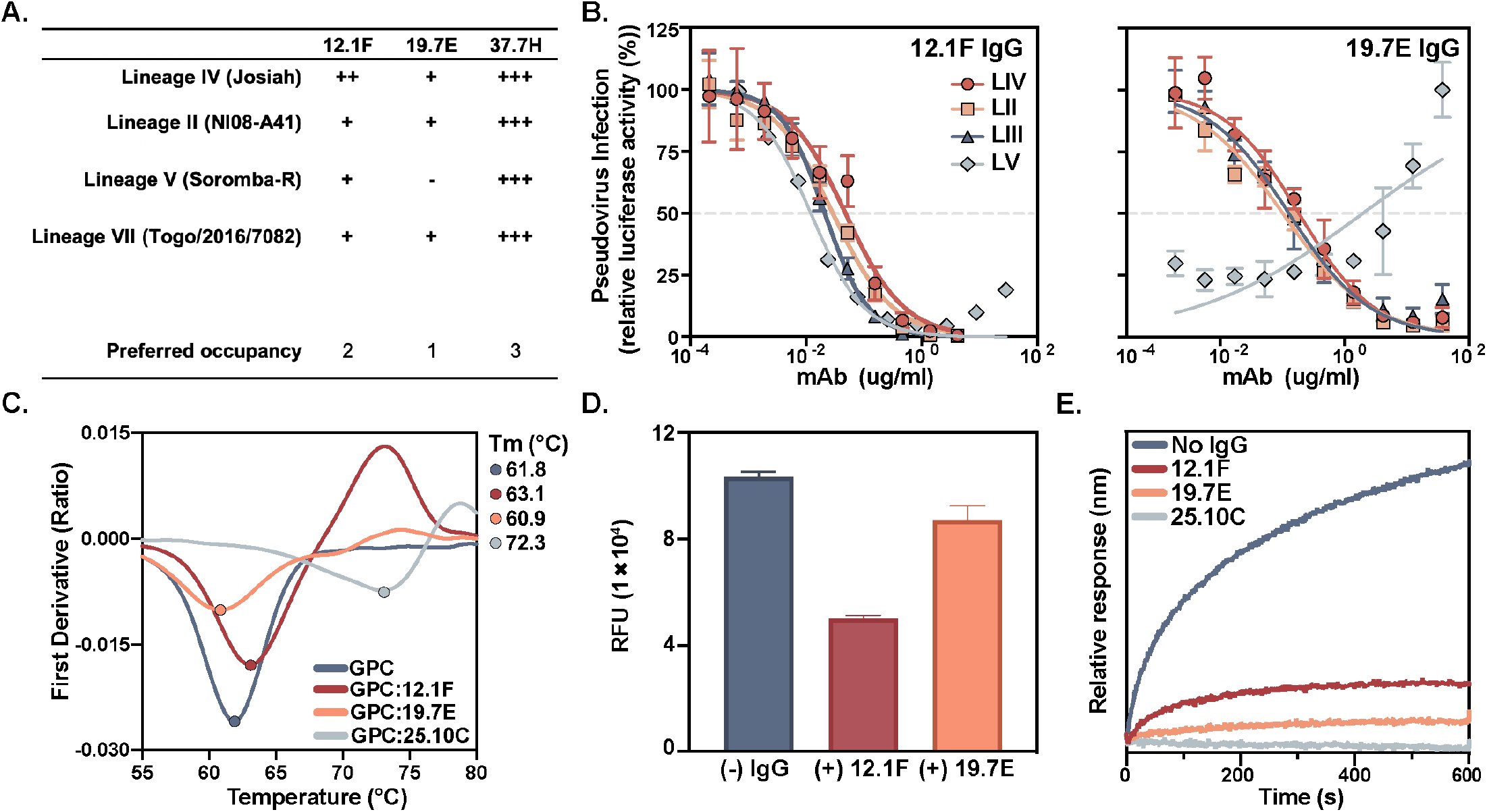
Characterization of the neutralizing GP1-A mAbs 12.1F and 19.7E. (A) Summary of mAb binding to GPCs by BLI (raw data in Fig. S6). The binding efficiency is based on the relative on-rate of IgG to immobilized GPC and is indicated as follows: +++, very strong binding; ++ strong binding; +, moderate binding; -, minimal to no binding. Proposed IgG occupancy per GPC is estimated based on relative R_max_ values under the assumption the highest R_max_ indicates full occupancy and 37.7H has a preferred occupancy of 3 Fabs per trimer (Hastie et al., 2019). (B) mAb neutralization of pseudoviruses derived from LASV strains of diverse lineages. The dotted line indicates 50% neutralization. Data points represent the mean with error bars indicating the SEM of three technical replicates (37.7H neutralization assay comparisons shown in Fig. S7A). (C) Thermostability of LASV LIV GPC-I53-50A in complex with indicated Fabs assessed by nanoDSF. Points represent the melting temperature (*T*_*m*_) of each complex. Each melting curve is a representative of triplicate curves with melting temperatures within ±0.1°C. (D) Synthetic matriglycan binding competition microarray of StrepTagged GPC-A binding to matriglycan with and without pre-treatment with 12.1F and 19.7E IgG. GPC-I53-50A bound to matriglycan was detected using StrepMAB antibody (Fig. S7E). (E) BLI competition analysis of immobilized GPC bound to indicated IgG and then exposed to recombinant LAMP1 at a pH of 5 (Fig. S7F).

While 12.1F maintained binding to all GPCs tested, 19.7E showed no binding to LV GPC and weaker relative binding to all other lineages. Both GP1-A mAbs demonstrated a benefit from avidity effects, with both 12.1F and 19.7E showing higher dissociation rates of Fabs compared to IgGs (Fig. S7A, B, and C). Furthermore, we were able to estimate the preferred binding stoichiometry of these Fabs based on the proportional R_max_ values relative to 37.7H (Fig. S6)—which is assumed to bind with one Fab per protomer based on previous work (i.e. three Fabs per trimer, Hastie et al., 2019). We showed 12.1F and 19.7E bind with lower preferred occupancies of two or one Fab per trimer, respectively, and this result was further corroborated by cryoEM (Fig. S7D). To assess differences in neutralization breadth, pseudovirus neutralization assays were performed. The 12.1F mAb was able to neutralize LASV lineage II, III, IV, and V while 19.7E neutralized II, III, and IV, but not V, consistent with the binding data. In line with our findings from BLI, 19.7E heavily relies on avidity for neutralization and is unable to neutralize LV virus (Fig. 3A, Fig. S6, and Fig. S7B and C). Curiously, 12.1F shows a strong reliance on avidity to neutralize LIII and LV virus yet had comparable potency against LIV and LII when used in IgG or Fab format.

To elucidate the mechanism of binding and neutralization for these mAbs, we performed nano differential scanning fluorimetry (nanoDSF) experiments, a matriglycan microarray competition assay (Sheikh et al., 2022), and BLI-based LAMP-1 competition experiments (Fig. 3C, D, and E). The GPC-B mAb 25.10C is known to stabilize the GPC’s prefusion conformation (Enriquez et al., 2022). Consistent with this finding, we observed 25.10C dramatically increased the melting temperature (*T*_m_) of LIV GPC-I53-50A by >10°C. In contrast, 12.1F and 19.7E had only marginal effects on GPC thermostability, suggesting these GP1-A mAbs likely do not neutralize by stabilizing the prefusion state of GPC. To probe the interaction between GPC and the matriglycan moieties of β-dystroglycan, we used a synthetic matriglycan microarray (Sheikh et al., 2022). This array presents chemoenzymatically-generated matriglycan oligosaccharides of defined length and shows length-dependent binding of LIV GPC-I53-50A to matriglycan, consistent with previous observations of GP1 and pseudovirus binding (Fig. S7E, Sheikh et al., 2022). Whereas LIV GPC-I53-50A showed strong binding to the microarray with 24 repeating disaccharide units, the same protein complexed with 12.1F bound markedly less. In contrast, 19.7E showed lower inhibition of matriglycan binding (Fig. 3D, Fig. S7E). Interestingly, the GPC-A mAb 25.10C also inhibited matriglycan binding while GPC-B mAb 37.7H did not (Fig. S7E). Furthermore, both mAbs show strong inhibition of GPC binding to recombinant LAMP-1 at pH 5 with inhibition levels comparable to 25.10C, which has been shown to completely block GPC binding to LAMP-1 (Fig 3E; Enriquez et al., 2022).

### Structural characterization of 12.1F and 19.7E mAbs reveals glycan dependence

To assess the molecular interactions of GP1-A antibodies to GPC, we used single-particle cryoEM and solved the structures of 12.1F and 19.7E bound to LIV GPC (Fig. 4A and B) to 3.7 and 3.8 Å, respectively (PDB 8EJH and 8EJI; EMDs 28182 and 28183). Our models reveal both antibodies bind near the apex of the trimer, with each Fab engaging a single GP1 subunit on the loop that extends over β5–β8. 12.1F uses both its HC and LC to interact with the GPC while 19.7E almost exclusively relies on its HC. 12.1F and 19.7E both bind in the space between apical glycans N89, N109, and N167, and show extensive contacts with the GPC glycans with total buried surface areas of 1549 and 1123 Å^2^, respectively.

**Fig 4:**
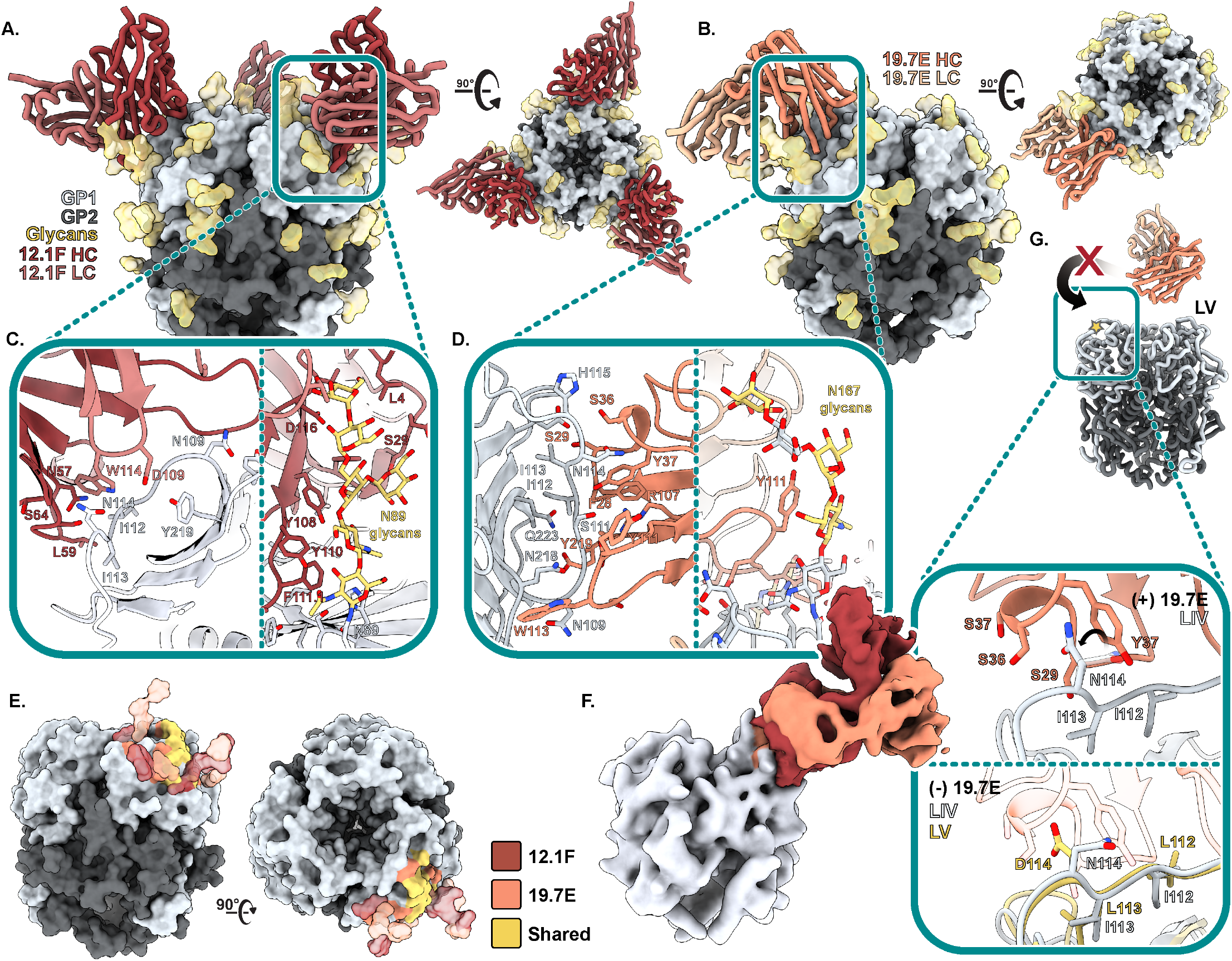
Structural description of the GP1-A epitope cluster. (A) Atomic model of LIV GPC (gray) bound to 12.1F Fab (red) determined by cryo-EM (PDB 8EJH, EMD-28182). (B) Atomic model of LIV GPC (gray) bound to 19.7E Fab (orange) determined by cryo-EM. (C) Key interactions between GP1 and 12.1F Fab at the epitope-paratope interface (PDB 8EJI, EMD-28183). Glycans within close proximity (<4 Å) shown in gold. More details can be found in Table S2. (D) Key interactions between GP1 and 19.7E Fab at the epitope-paratope interface. Glycans within close proximity (<4 Å) shown in gold More details can be found in Table S3. (E) The GP1-A antigenic landscape mapped on LIV GPC and colored according to the 12.1F (red), 19.7E (orange), or shared (yellow) antibody footprint. Glycan contacts are noted as transparent surfaces coloured according to Fab interaction. (F) Overlaid, gaussian-filtered maps showing the angle of approach taken by 12.1F (red) and 19.7E (orange) Fabs to engage LIV GPC. (G) Analysis of the residues at the 19.7E binding site for LIV and LV GPCs. The gold star indicates the loop in the panels below. The top panel shows the LIV GPC conformation when bound to 19.7E. Residues I112, I113 and N114 are shown as their mutation in LV (I112L, I113L, and N114D, Fig. S1A) prohibits 19.7E binding. The rotameric shift of LIV N114 in the 19.7E-bound state is indicated. The bottom panel shows a comparison of unbound LIV and LV GPCs with 19.7E shown in translucent orange to indicate its positioning when bound to LIV GPC. marked residues indicate differences in the amino acid sequences between LIV and LV.

While our previous observations and 2D classifications suggest 12.1F typically binds in a 2 Fab per 1 GPC fashion, applying C3 symmetry to the data enabled the best resolution of the epitope-paratope interaction (Fig. 4C). Amino acid residues at the epitope-paratope site primarily interact through hydrogen bonding with the residues past the β5 sheet and before the apex-associated α1 helix with the HC’s CDRH2 loop providing the most notable amino acid contacts (N57, L59, S64, and T65; Fig 1B, left; IMGT numbering). The LC’s CDRL3 residues predominantly engage with GP1’s S111 to contribute additional stability through hydrogen bonding. The CDRL2 sits beside the N89 glycan. The 18 amino acid CDRH3 loop of 12.1F, while in close proximity (<4 Å) to GP1 residues, only weakly associates with GP1 amino acids. Instead, the CDRH3 makes extensive contacts with the apex glycans despite the small variability observed in Fig 2A. The HC interacts heavily with the N89 glycans and multiple aromatic residues (Y108, Y110, F111.1; IMGT numbering) engage with the sugar moieties (Fig. 4C, right). This trend extends to glycan N109, which interacts with W112 of the HC. The glycans we modeled contribute 59% of the total buried surface area between the Fab and GPC with individual glycan contributions of 547 Å^2^ (N89), 191 Å^2^ (N109), and 33 Å^2^ (N167). Additional contacts are described in Table S2. We observed density for the fusion peptide of GPC bound to 12.1F in two conformations: (1) similar to unbound GPCs (Fig. 2E) and (2) similar to 18.5C, 37.7H, 25.6A, and 25.10C antibodies (Fig. 2E, Fig. S2B).

For 19.7E, we typically only saw one Fab bound per GPC trimer and symmetry-expanded particles to achieve a subset of protomers bound to the Fab. This antibody makes more contacts with amino acid residues than 12.1F (Fig. 4D, left; Table S3), entirely via the HC. 19.7E engages with GP1 residues along the β-sheet surface using its CDRH1 and CDRH3 loops. Amino acid contacts of interest include GP1’s S111, which likely hydrogen bond with HC’s Y37, R107, and/or D112. Residues I112 and I113 also have multiple potential hydrogen bonding partners including S29 and Y37. While most interactions at this interface are facilitated by hydrogen bonding, hydrophobic packing between GPC’s Y172 and the W113 of the CDRH3 loop as well as GPC’s I112 with F28 and 2V of the HC also contribute to the antibody’s ability to bind GPC. While 19.7E also utilizes the apex N89, N109, and N167 glycans (Fig. 4D, right), it shares considerably fewer interacting partners when compared to 12.1F (Tables S2 and S3). The LC only interacts minimally with the N89 and N109 glycans. The modeled GPC glycans contribute 47% of the total buried surface area when 19.7E binds to GPC with individual glycan contributions of 251 Å^2^ (N167), 147 Å^2^ (N109), and 128 Å^2^ (N89). Upon GPC binding to 19.7E, the fusion peptide takes on a similar conformation as seen with GPC-A and GPC-B NAbs and extends toward the trimer interior (Enriquez et al., 2022; Hastie et al., 2017, 2019).

As we noticed the GP1-A antibodies shared extensive interaction networks with the apex glycans, we decided to assess whether neutralization by these mAbs is glycan-dependent, as has been seen previously with the NAb LAVA01 (Brouwer et al., 2022). While we observed exceptional interactions of both NAbs with the N89 glycan, previous studies indicate N89 glycan removal leads to cleavage inefficiency. Similarly, an N109Q or N109A substitution also leads to reduced proteolytical processing (Zhu et al., 2021). Therefore, we generated pseudoviruses containing the S111A and N167Q glycan knockout mutations. The 12.1F mAb’s neutralization potency was drastically reduced after knocking out the N109 glycan. The 19.7E mAb required both the N109 and N167 glycans to neutralize LIV LASV pseudovirus (Fig. S8A).

Inspection of the structures support the LAMP-1 and matriglycan competition we observed for these GP-1 mAbs. The 12.1F and 19.7E Fabs come within close proximity of H92 (Table S2), which—together with H93 and H230—constitutes the histidine triad and regulates onset of pH-dependent conformational changes in GP1 required for LAMP-1 binding (Acciani et al., 2017; Cohen-Dvashi et al., 2016; Israeli et al., 2017). While there are no additional contacts between 12.1F and 19.7E and the putative LAMP-1 binding site outside of H92 (Fig. S8B), it is likely the Fabs are inhibiting LAMP-1 binding through steric hindrance or by disabling the required conformational changes. We observed an apparent discrepancy when inspecting the location of the 12.1F and 19.7E epitopes and the extent of matriglycan competition. Whereas 12.1F showed a much stronger ability to compete with matriglycan than 19.7E, the latter makes closer molecular contacts to the apex of GPC (Fig. 4E). Regardless, the interactions at both epitope-paratope interfaces do not directly interfere with residues known to associate with matriglycan (Fig. S8C, Katz et al., 2022). The results can be reconciled by considering the angles of approach of these mAbs as we observed 12.1F Fab engaged at a steeper angle relative to the GPC’s three-fold symmetry axis, which presumably causes steric impediment of matriglycan engagement. (Fig. 4F).

Our structures (Fig. 2), enable mapping of single point mutations responsible for antigenic differences among LASV lineages and analysis of accompanying structural ramifications. We observe that mutations at residues 112-114 are likely to be responsible for the loss of 19.7E neutralization against LV. An overlay of the structures of unliganded LIV GPC with that of LIV in complex with 19.7E shows N114 adjusting its rotameric position upon Fab binding and positions itself among three serine residues of the CDRH1 (Fig. 4G, top). Comparison of unliganded and bound LIV GPC shows binding of 19.7E displaces the 112-114 residues by an average of 1.3 Å. In the unbound state, LV residues 112-114 (Fig. 4G, gold star) extend further away from the b-sheet surface and would need to be displaced by an average of 2.1 Å to adopt the same conformation. Additionally, the D114 of LV GPC likely disrupts the electrostatic complementarity of N114 with the surrounding S29, S36, and S37 residues resulting in the escape of LV to 19.7E (Fig. 4G, bottom).

### Novel mAb S370.7 binds to the GPC-B epitope cluster and prefers trimer over monomer

We previously showed GPC-I53-50A proteins represent useful baits for antigen-specific B cell sorting (Brouwer et al., 2022). To expand the repertoire of available anti-GPC mAbs we used LIV GPC-I53-50A as a bait for antigen-specific B cell sorting of convalescent serum from patient 1102370, a member of the Lassa fever survivor cohort at the Kenema Government Hospital (Robinson et al., 2016). In doing so, we isolated a novel antibody, S370.7, which binds with high affinity to GPC (Fig. 5A) and neutralizes LIV pseudovirus with an IC_50_ of 0.45 μg/mL (Fig. 5B). Similar to the GP1-A antibodies, S370.7 benefited from avidity, as evidenced by the increased off-rate of Fab from GPC compared to IgG (Fig. 5C). S370.7 only marginally increased the stability of the GPC-I53-50A trimer by nanoDSF in contrast to antibodies 25.10C and 37.7H (Fig. 5D, Fig. S9A). Interestingly, this mAb does not inhibit LAMP-1 binding nor block matriglycan attachment to the GPC (Fig. 5E and F).

**Fig 5:**
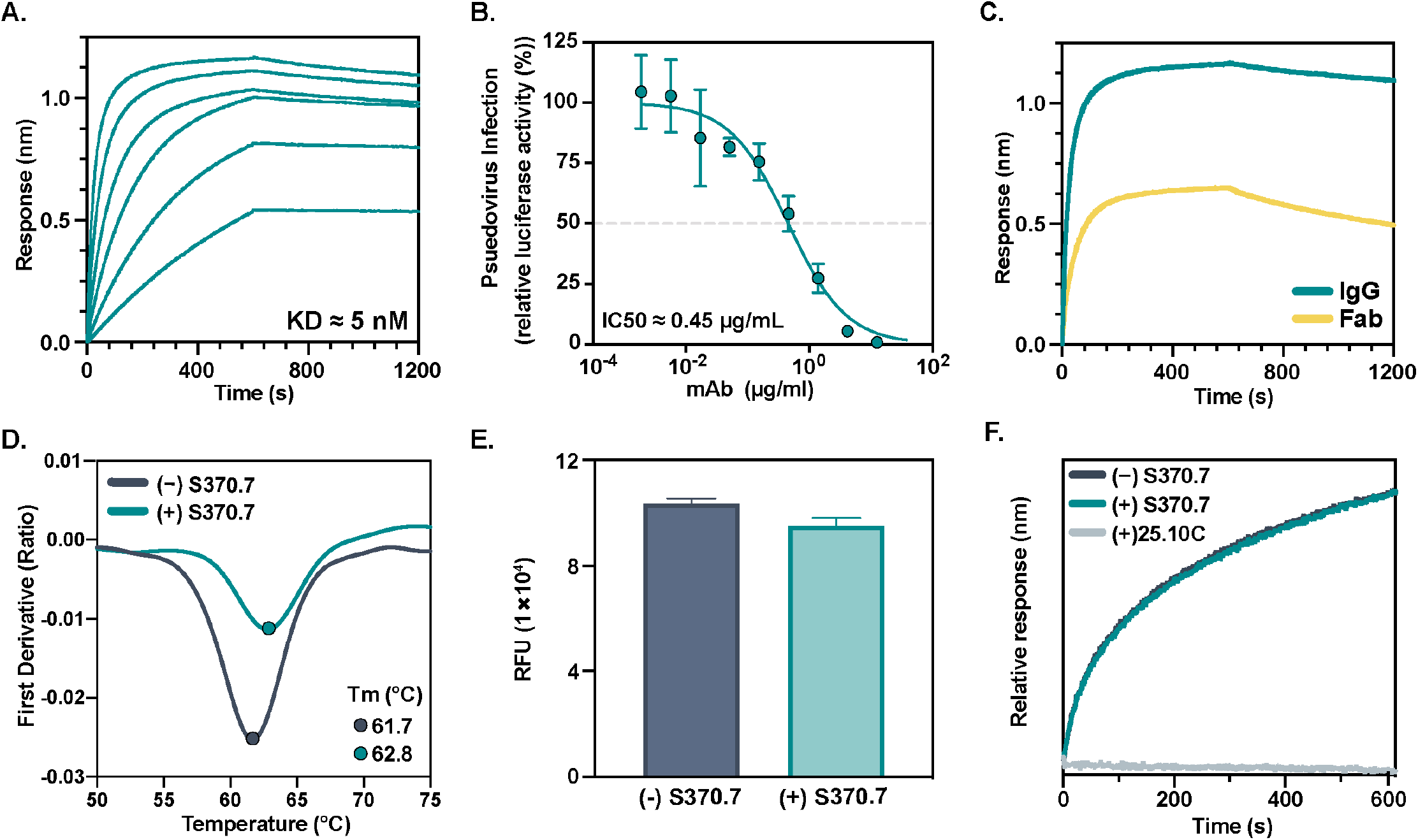
Isolation of a novel monoclonal NAb using GPC-I53-50A. (A) BLI sensorgrams depicting immobilized GPC-I53-50A binding to S370.7 IgG in a dose-dependent manner. IgG concentrations used were 400, 200, 100, 50, 25, and 12.5 nM. K_D_ value determined using a 1:1 binding profile and assuming partial dissociation. (B) LIV LASV pseudovirus neutralization of LASV by S370.7. The dotted line indicates 50% neutralization. Data points represent the mean with error bars indicating the SEM of three technical replicates. (C) BLI sensorgram comparing binding of S370.7 IgG to Fab to immobilized GPC. IgG and Fab were added at an equimolar concentration of 400 nM. (D) Thermostability of LIV GPC in complex with S370.7 assessed by nanoDSF. Points represent the melting temperature (T_m_). Each melting curve is a representative of triplicate curves with melting temperatures within ±0.1°C. (E) Synthetic matriglycan binding microarray of StrepTagged GPC-I53-50A bound to S370.7 IgG and detected using StrepMAB antibody. (F) BLI analysis of immobilized GPC bound to S370.7 or 25.10C IgG and then exposed to recombinant LAMP-1 at a pH of 5.

To further probe the molecular interactions between S370.7 and GPC, we solved a 3.2 Å structure of GPC-I53-50A bound to S370.7 Fab by single-particle cryoEM (Fig. 6A). The model reveals S370.7 engages two adjacent protomers of the GPC with interactions almost exclusively within GP2. S370.7 HC and LC primarily contact separate protomers of the GPC. The HC, which features a longer, 22 amino acid CDRH3 loop than is typical for anti-LASV antibodies (Robinson et al., 2016) and has a 6.5% somatic hypermutation in its IGHV4-34*02 gene, penetrates the pocket situated between the fusion peptide of one protomer and the HR1d, HR2, and T-loop domains of the neighboring protomer. Both HC and LC are flanked by the N390 and N79 glycan, respectively, with minor contacts made between each (Table S3).

**Fig 6:**
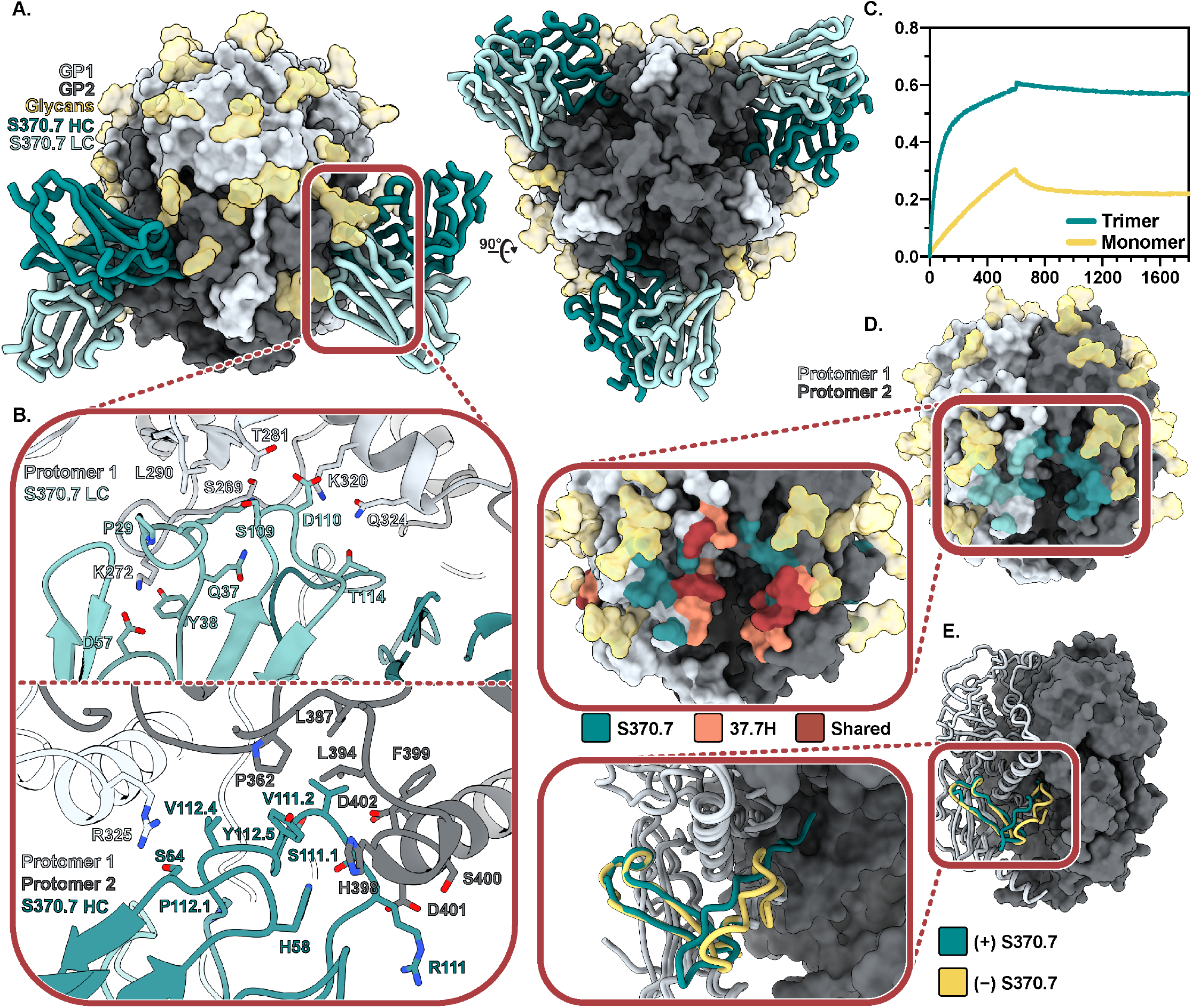
Structural characterization of the trimer-preferring NAb S370.7. (A) Atomic model of LIV GPC (gray) bound to S370.7 Fab (teal) determined by cryo-EM (PDB 8EJJ, EMD-28184). (B) Key interactions between S370.7 LC (top) and HC (bottom) residues with GPC. More detailed information can be found in Table S4. (C) BLI sensorgram showing the binding profile of immobilized S370.7 IgG to GPC trimer or GPC monomer in equal protomer concentrations. (D) S370.7 antibody footprint. HC interactions are shown in dark teal and LC interactions in light teal. Pop-out image shows the overlap and distinctions with known GPC-B NAb 37.7H. (E) Comparison of the fusion peptides of S370.7-bound LIV GPC (teal) with unbound LIV GPC (PDB 8EJD; yellow).

The LC, which features a 3.8% somatic hypermutation rate in its IGLV3-25*03 gene, engages the GPC (Fig 6B, top) with its CDRL1 (11 amino acids), CDRL2 (8 amino acids), and CDRL3 loops (11 amino acids). The CDRL1 forms hydrogen bonds with residues K272 and S269 of the GP2 fusion peptide. Additionally, K272 interacts with the CRL2 loop as well, forming a salt bridge with D57. The CDRL3 loop residues D110 and T114 form hydrogen bonds with K320 and Q324 of the HR1 helix. D110 and K320 likely engage further and form a salt bridge, strengthening the interaction. While the HC almost exclusively interacts with GPC via its CDRH3 (Fig 6B, bottom), its 7 amino acid CDRH2 putatively forms a hydrogen bond at S64 with the R325 of GPC, making it the strongest cross-protomer interaction of the HC. Interactions between hydrophobic residues of the CDRH3 (Y112.5, V112.4, and V111.2) with residues just upstream of and extending to the HR2 helix of GP2 (L387, S389, L394, F399) support antibody binding by forming a stable, hydrophobic pocket. Just beyond this hydrophobic pocket, there appears to be an additional favorable electrostatic interaction forming between D401 of GP2 and R111 of the CDRH3. The total buried surface area between the Fab and GPC is 1275 Å^2^ of which the modeled glycans contribute 22%.

Based on the nature of the S370.7 epitope, we hypothesized S370.7 would require the correct quaternary presentation of GPC to bind. We found that while S370.7 could still bind LIV GP monomer, it did so at a reduced rate with higher dissociation compared to its binding to LIV GPC trimer, making the S370.7 trimer-preferring. Compared to other known antibodies (Fig. S9B), S370.7 exhibits the highest degree of preference for the trimeric conformation of GPC. We compared the epitopes of S370.7 and 37.7H and observe a marked overlap, especially within the region upstream of the HR2 helix, making S370.7 a member of the GPC-B competition group of anti-LASV mAbs (Fig. 6D). We also note a conformational change in the fusion peptide upon binding to S370.7 (Fig. 6E), that is consistent with our observations of other antibody-bound fusion peptide conformational differences (Fig. 2E and Fig. S2B). Its lack of matriglycan inhibition and dissociation from GPC at a pH of 5.0 (Fig. S7F) suggest anti-LASV GPC antibodies can exhibit alternative neutralization mechanisms that have yet to be elucidated.

## Discussion

The advancement of prefusion-stabilized GPCs is an important step for developing useful immunogens capable of overcoming the notoriously poor humoral immune response to LASV and LASV vaccines (Fisher-Hoch et al., 1989; McCormick and Fisher-Hoch, 2002). Here, we further demonstrate the use of the I53-50A protein as a trimerization scaffold for the stabilization of GPCs of four of the seven LASV lineages, three of which we describe structurally for the first time. The GPC-I53-50As are a suite of stable, soluble heterologous proteins useful for assessing cross-binding and are amenable to cryoEM analysis when high-resolution information is needed. Importantly, the GPC-I53-50As present native-like epitopes and bind to antibodies within the canonical GP1-A, GPC-A, and GPC-B competition groups without the need for additional stabilizing antibodies. Our unliganded GPCs enable tracking of the fusion peptide response to antibody binding, thus enabling more complete insights into the binding and neutralization mechanisms of anti-GPC antibodies.

The GPC structures bound to 12.1F and 19.7E presented here define the GP1-A competition group and show their epitope resides near the apex of the GP1 protomer and interacts widely with apical glycans. Glycan-dependence is confirmed through complementary glycan knockout pseudovirus neutralization assays. These antibodies contribute to LASV neutralization by hindering GPC’s ability to (1) bind the matriglycan sugars of its extracellular receptor α-dystroglycan and (2) engage with the endosomal receptor LAMP-1. Intriguingly, we also observed GPC-A mAb 25.10C inhibits matriglycan binding. The inhibition of GPC binding to both matriglycan and LAMP-1 by 12.1F and 25.10C mAbs may explain their more potent neutralizing properties compared to other isolated mAbs especially in light of findings that LAMP-1 is not necessary for LASV fusion (Hulseberg et al., 2018; Markosyan et al., 2021; Robinson et al., 2016; Zhang et al., 2022).

Finally, we demonstrate GPC-I53-50As are valuable baits for antigen-specific B cell sorting with our discovery of GPC-B NAb S370.7 using LIV GPC-I53-50A. This novel antibody engages the GPC in a similar fashion as the majority of neutralizing anti-GPC antibodies (Robinson et al., 2016) and uses both its HC and LC, which are flanked by N79 and N390 glycans, to engage adjacent protomers. Binding by S370.7 causes migration of the fusion peptide to the interior of the trimer, where it resides beneath the C-terminal of GP2. Intriguingly, we were not able to confirm the neutralization mechanism of S370.7, which likely indicates this mAb acts through an unknown neutralization mechanism or points to a limitation of pseudovirus assays for the Old World arenaviral family.

Using additional GPC-I53-50A lineages as probes, we can now sort B-cells with broader LASV specificity for the discovery of novel mAb therapeutics. Further, applying this stabilization scheme to additional arenaviruses presents the exciting opportunity to screen for antibodies capable of binding across Old World and New World arenaviruses. In summary, our findings and the suite of GPC ectodomains (1) informs more comprehensive immunogen design and stabilization work, specifically in the context of GP1-A antibodies (2) describes stable, trimeric GPC reagents for cross-neutralization assessment, and (3) provides a robust and relatively high-throughput platform for single-particle cryoEM analysis of LASV GPCs with and without NAbs.

## Methods

### Sequence alignment and conservation assessment

S genomes of LASV field isolates (Li and Tian, 2020) were aligned, matched to groups according to codon reading frame, and re-aligned based on amino acid residue using Clustal Omega multiple sequence alignment (Sievers and Higgins, 2018). A total of 361 GPC sequences were analyzed. Conservation is visualized using AL2CO (Pei and Grishin, 2001) entropy measure with the modified Henikoff & Henikoff frequency estimation method and a gap fraction of 0.7 and visualized in ChimeraX (Pettersen et al., 2021).

### Construct design

The LIV GPC monomer, LIV GPC-I53-50A, and Avi-his-tagged LIV GPC-I53-50A constructs were generated as described previously (Brouwer et al., 2022). To generate the NIG08-A41-, Soromba-R, and Togo/2016/7082-GPC-I53-50A constructs, genes encoding GPC residues 1-423 (Genbank: ADU56626.1), 1-424 (Genbank AHC95553.1), and 1-423 (Genbank AMR44577.1), respectively, with the GPCysR4 mutations introduced (Hastie et al., 2017) were cloned by Gibson assembly into PstI-BamHI-digested Josiah-GPC-I53-50A plasmid. A LIV GPC-I53-50A construct with the native site-1 protease cleavage site was generated by introducing R258L and R259L mutations by Q5 site-directed mutagenesis. The 12.1F, 19.7E, 37.7H, and 25.10C sequences were derived from patent WIPO: WO2018106712A1. The 19.7E, 37.7H, 12.1F, 25.10C, and S370.7 plasmids were generated by Gibson assembly of genes encoding the variable regions of the corresponding heavy and light chains into plasmids containing the constant regions of the human IgG1 for the heavy or light chain. Plasmids encoding histidine-tagged Fab regions of 12.1F, S370.7, and 25.10C were generated by introducing a histidine-tag followed by a stop-codon in the hinge region (directly upstream of the DKTHT motif) of the corresponding heavy chain plasmid by Q5 site-directed mutagenesis. For pseudovirus neutralization assays, a pPPI4 plasmid was digested with PstI-NotI and a gene encoding full-length GPC of lineage II (NIG08-A41), lineage III (CSF; Genbank: AAL13212.1), or lineage V (Soromba) was inserted by Gibson assembly. Q5 site-directed mutagenesis was used to introduce the S111A and N167Q mutations into a plasmid encoding full-length Josiah GPC (a kind gift from Robin Shattock).

### Protein expression and purification

GPC-I53-50As, LIV GPC monomer, biotinylated GPC-I53-50As and antibodies were transiently expressed in HEK 293F cells at a density of 1.0 × 10^6^ cells/mL using PEImax at a ratio of 1:3 DNA to PEI. HEK 293F cells were maintained in HEK 293F in 293FreeStyle expression medium (Life Technologies) and cultured at 37°C with 8% CO_2_ while shaking at 125 rpm. Plasmids encoding GPCs were co-transfected with a furin plasmid at a 1:2 ratio. To express biotinylated GPC-I53-50A, HEK 293F cells were co-transfected with Avi-his-tagged GPC-I53-50A, furin and a BirA plasmid (a kind gift from Lars Hangartner) in a 2:1:0.5 ratio. IgG plasmids were transfected at a heavy and light chain ratio of 1:1 while the generation of Fabs of 12.1F, 25.10C, and S370.7 was initiated by transfecting the histidine-tagged heavy chain Fab domain with the corresponding light chain at a ratio of 1:2. Culture supernatants of GPC constructs were harvested after six days, while IgG and Fab were harvested after five days. GPC-I53-50As were purified by gravity column using StrepTactin 4Flow resin (IBA Life Sciences) according to manufacturer’s protocol and eluted with 1X BXT (IBA Life Sciences). IgGs were purified by gravity column using Protein G resin (Cytiva) and eluted with 0.1 M glycine at pH 2.0. Biotinylated GPC-I53-50As and Fabs of 12.1F and S370.7 were purified by rolling the culture supernatant overnight at 4°C with Ni-NTA Agarose resin (Thermo Scientific). The next day, the bead suspension was flown over a gravity column, washed with 20mM imidazole, 50 mM NaCl, pH 7.0 and eluted with 500 mM imidazole, 50 mM NaCl buffer, pH 7.0. Recombinant LAMP-1 was generated by transfecting HEK 293F cells with a rabbit Fc-tagged LAMP-1 plasmid encoding residues A29-S351 (a kind gift from Thijn Brummelkamp, Jae et al., 2014). Culture supernatant was then incubated with CaptureSelect IgG-Fc resin (Thermo Scientific) and eluted from the resin using 0.1 M glycine, pH 3.0, into neutralization buffer (1 M Tris, pH 8.0) at a 1:9 ratio. All proteins were buffer exchanged to TBS after elution and purified further by size exclusion chromatography using a Superdex 200 increase 10/300 GL column (Sigma-Aldrich) with TBS as its running buffer. Fractions corresponding to the appropriate peaks were concentrated using a MWCO concentrator with the following cutoffs: 100 kDa for GPC-I53-50As; 30 kDa for IgGs and LIV GPC monomer; and 10 kDa for Fabs (Millipore).

### Differential scanning fluorimetry

Thermostability of GPC and GPC-Fab complexes was determined with a nano-DSF NT.48 (Prometheus). GPC proteins or complexes were diluted to 0.5 mg/mL and loaded into high sensitivity capillaries. The assay was run with a linear scan rate of 1°C/min and 80%-100% excitation power. The first derivative of the ratio of tryptophan fluorescence wavelength emissions at 350 and 330 nM were analyzed to determine thermal onset (*T*_*onset*_) and denaturation (*T*_*m*_) temperatures using the Prometheus NT software.

### Negative stain electron microscopy

Carbon-coated 400-mesh copper grids were glow discharged for 25 s at 15 mA using a PELCO easiGlow instrument (Ted Pella, Inc.). GPC-I53-50A samples were diluted in TBS to approximately 15 μg/mL and loaded onto the copper grids and incubated for 30 s. The sample was blotted and immediately stained with 2% (w/v) uranyl formate for 15 s. Excess stain was removed by blotting and grids were dried for >5 minutes before being loaded on a 200 kV Tecnai F20 electron microscope (FEI) featuring a TemCam F416 CMOS camera (TVIPS). Images were collected at a magnification of 62,000X with a defocus value of -1.5 um, total electron dose of 25 e^-^/Å^2^, and pixel size of 1.77 Å. Images were acquired using the Leginon software package (Suloway et al., 2005). Approximately 100,000 particles were picked using Appion (Lander et al., 2009) and 2D classification was performed with Relion 3.0 (Zivanov et al., 2018).

### Site-specific glycan analysis

100 μg aliquots of each sample were denatured for 1h in 50 mM Tris/HCl, pH 8.0 containing 6 M of urea and 5 mM dithiothreitol (DTT). Next, GPC-I53-50A samples were reduced and alkylated by adding 20 mM iodoacetamide (IAA) and incubated for 1h in the dark, followed by a 1h incubation with 20 mM DTT to eliminate residual IAA. The alkylated GPC-I53-50A samples were buffer exchanged into 50 mM Tris/HCl, pH 8.0 using Vivaspin columns (3 kDa) and two of the aliquots were digested separately overnight using chymotrypsin (Mass Spectrometry Grade, Promega) or alpha lytic protease (Sigma Aldrich) at a ratio of 1:30 (w/w). The next day, the peptides were dried and extracted using C18 Zip-tip (MerckMilipore). The peptides were dried again, re-suspended in 0.1% formic acid and analyzed by nanoLC-ESI MS with an Ultimate 3000 HPLC (Thermo Fisher Scientific) system coupled to an Orbitrap Eclipse mass spectrometer (Thermo Fisher Scientific) using stepped higher energy collision-induced dissociation (HCD) fragmentation. Peptides were separated using an EasySpray PepMap RSLC C18 column (75 μm × 75 cm). A trapping column (PepMap 100 C18 3 μM 75 μM x 2cm) was used in line with the LC prior to separation with the analytical column. The LC conditions were as follows: 280 minute linear gradient consisting of 4-32% acetonitrile in 0.1% formic acid over 260 minutes followed by 20 minutes of alternating 76% acetonitrile in 0.1% formic acid and 4% ACN in 0.1% formic acid, used to ensure all the sample had eluted from the column. The flow rate was set to 200 nL/min. The spray voltage was set to 2.7 kV and the temperature of the heated capillary was set to 40°C. The ion transfer tube temperature was set to 275°C. The scan range was 375–1500 m/z. Stepped HCD collision energy was set to 15, 25 and 45% and the MS2 for each energy was combined. Precursor and fragment detection were performed using an Orbitrap at a resolution MS1= 120,000. MS2= 30,000. The AGC target for MS1 was set to standard and injection time set to auto which involves the system setting the two parameters to maximize sensitivity while maintaining cycle time. Full LC and MS methodology can be extracted from the appropriate raw file using XCalibur FreeStyle software or upon request.

Glycopeptide fragmentation data were extracted from the raw file using Byos (Version 4.0; Protein Metrics Inc.). The glycopeptide fragmentation data were evaluated manually for each glycopeptide; the peptide was scored as true-positive when the correct b and y fragment ions were observed along with oxonium ions corresponding to the glycan identified. The MS data was searched using the Protein Metrics 305 N-glycan library with sulfated glycans added manually. The relative amounts of each glycan at each site as well as the unoccupied proportion were determined by comparing the extracted chromatographic areas for different glycotypes with an identical peptide sequence. All charge states for a single glycopeptide were summed. The precursor mass tolerance was set at 4 ppm and 10 ppm for fragments. A 1% false discovery rate (FDR) was applied. The relative amounts of each glycan at each site as well as the unoccupied proportion were determined by comparing the extracted ion chromatographic areas for different glycopeptides with an identical peptide sequence. Glycans were categorized according to the composition detected.

### CryoEM grid preparation and imaging

To prepare grids for sample application, UltrAuFoil R1.2/1.3 (Au, 300-mesh; Quantifoil Micro Tools GmbH) grids were treated with Ar/O^2^ plasma using a Solarus plasma cleaner (Gatan) for 10 s or were plasma discharged for 25 s at 15 mA using a PELCO easiGlow (Ted Pella Inc.). Right before applying the protein samples to the grids, we added flouro-octyl maltoside at a final concentration of 0.02% (w/v). Cryo-grids were prepared using a Vitrobot mark IV (Thermo Fisher Scientific). In all instances, the chamber temperature and humidity were set to 4°C and 100%, respectively. Samples were frozen using variable blot times between 3 to 7 s with a blot force of 1 s and a wait time of 10 s. After blotting, the grids were plunge-frozen in liquid ethane.

Cryo-grids were loaded into an FEI Titan Krios or Talos Arctica (Thermo Fisher Scientific), which operate at 300 or 200 kV, respectively. Both microscopes were equipped with a K2 Summit direct electron detector camera (Gatan). The data were collected with approximate cumulative exposure of 50 e^-^/A^2^. Magnifications were set to 130,000 or 36,000X for the Krios and Arctica, respectively. Automated data collection using the Leginon software package (Suloway et al., 2005) was employed for all datasets reported. Additional information can be found in Table S1.

### CryoEM data processing

Image preprocessing was performed using the Appion software package (Lander et al., 2009). Micrograph movie frames were first aligned and dose-weighted using the UCSF MotionCor2 software (Zheng et al., 2017b). Initial data processing was performed in cryoSPARC v3.0 (Punjani et al., 2017) including particle picking and early 2D classification. Quality initial 2D classes were used to inform template picking of the datasets followed by iterative rounds of 2D classification where bad particle picks were removed.

All datasets were analyzed using an initial model generated in UCSF chimera (Pettersen et al., 2004) from known structures of the LIV GPC (PDB 7SGD) and I53-50A protein (PDB 6P6F). For GPC-I53-50A and Fab complexes, the ligand-free initial model was used for initial 3D refinement steps. After preliminary 3D maps were generated demonstrating Fab density, they were lowpass filtered and used as the initial model for subsequent steps.

For ligand-free GPC-I53-50As, preliminary 3D refinements were performed in cryoSPARC v3.0 (Punjani et al., 2017). Heterogeneous refinements were used to sort out remaining bad particles and homogenous refinements to orient the GPC appropriately by including the I53-50A scaffold density. Iterative rounds of local refinements were performed with masks that excluded the scaffold density. These particle stacks were transferred to Relion 3.1 (Zivanov et al., 2018) for further processing. Local 3D refinements and 3D classifications without global alignment were performed to further polish the particle stack. C3 symmetry was then applied during local 3D refinement followed by CTF refinements. Particle stacks were imported back to cryoSPARC v3.0 for final rounds of C3 local refinement, global CTF refinement, and the final C3 local refinement job. See Fig. S3B for more detail.

For antibody-bound GPC-I53-50A structures, the same general processing steps were followed as above sans moving particles to Relion 3.1. LIV GPC-I53-50As bound to 12.1F and S370.7 were analyzed by imposing C3 symmetry after initial alignments. LIV GPC-I53-50A bound to 19.7E was analyzed by symmetry expanding the particle set after C1 alignment along the C3 axis of symmetry. Particles were sorted using focused classification using a 60 Å sphere mask around the epitope-paratope interface to distinguish particles with Fab density. Subsequent refinements were performed to constrain particle alignment to one protomer face.

### Atomic model building and refinement

Post-processed maps were used to build all final atomic models. For LIV GPC-I53-50As, PDB 7SGD was used as the initial model and manually fit into density using Coot (Emsley and Crispin, 2018). Initial models for LII, LV, and LVI GPC-I53-50As were generated using SwissModeler (Waterhouse et al., 2018) and manually fit into density using Coot. 12.1F, 19.7E, and S370.7 Fab initial models were produced by ABodyBuilder (Leem et al., 2016) and manually fit into the post-processed maps using Coot (Emsley et al., 2010). Iterative manual modeling building in Coot followed by Rosetta relaxed refinement were used to generate the final models (Wang et al., 2016). The model fit to map for all models was validated using MolProbity and EMRinger analyses (Barad et al., 2015; Chen et al., 2010) in the Phenix software package (Liebschner et al., 2019). Epitope-paratope interactions were analyzed in UCSF ChimeraX (Pettersen et al., 2021) and the web-based Epitope-Analyzer (Montiel-Garcia et al., 2022). Buried surface area calculations for the fusion peptide and RMSD calculations were performed using UCSF Chimera (Pettersen et al., 2004). Buried surface area calculations for antibody interactions were calculated using PDBePISA (Krissinel and Henrick, 2007). Final atomic models have been submitted to the Protein Data Bank (PDB) with accession codes found in Table S1. All figures featuring atomic models were generated using UCSF ChimeraX (Pettersen et al., 2021).

### Antibody digestion and Fab purification

Fabs of 19.7E were generated by papain digestion of purified IgG. First, a buffered aqueous suspension of papaya latex papain (Sigma Aldrich) was activated by incubating in 100 mM Tris, 2 mM EDTA, 10 mM L-cysteine at 37°C for 15 mins. Next, IgG was incubated with activated papain in 100 mM Tris, 2 mM EDTA, 10 mM L-cysteine at a ratio of 40 μg activated papain per 1 mg of purified IgG for 5 hours at 37°C. The reaction was quenched by adding iodoacetamide to a final concentration of 0.03 M. Undigested IgG and Fc fragments were removed by a 2 h incubation with CaptureSelect IgG-Fc resin (Thermo Fisher Scientific). Resin was spun down and the supernatant run on a Superdex 200 increase 10/300 GL column (Sigma-Aldrich) size exclusion column using TBS as its running buffer. Fractions from 15.5-16.5 mL elution volume were collected and concentrated in a MWCO concentrator (Millipore) with a 10 kDa cutoff.

### Antibody affinity measurements using BLI

Antibody binding to GPC-I53-50As was assessed using an Octet Red96 instrument (ForteBio). Biotinylated GPC-I53-50A was loaded onto SA sensors (Sartorius) at 100 nM. After a short dip in running buffer (PBS, 0.1% BSA, 0.02% Tween20, pH 7.4), sensors were dipped in IgGs diluted to 400, 200, 100, 50, 25, or 12.5 nM. For Fab measurements the sensors were dipped in a 400 nM dilution of Fabs. Association and dissociation steps were measured for 600 s. Assays were performed at 30°C. All dilutions were made in running buffer with a final volume of 200 μL per well. 12.1F and 19.7E IgG kinetics were modeled assuming a 1:1 binding model while 37.7H assumed a 2:1 binding model.

### LAMP-1 competition assessment using BLI

Biotinylated GPC-I53-50As diluted in running buffer (PBS, 0.02% Tween20, 0.1% BSA) were loaded onto SA sensors (Sartorius) to a signal of 1.0 nm using an Octet Red96 system (ForteBio). After a short dip in running buffer, the sensors were dipped in 400 nM of 12.1F, 19.7E, S370.7, or 25.10C diluted in running buffer or running buffer alone. To measure IgG dissociation, the sensor was dipped for 1200 s in pH 5.0 running buffer (50 mM NaCitrate, 150 mM NaCl, pH 5.0, 0.1% BSA, 0.02% Tween20). The sensor was then dipped for 600 s in 200 μg/mL of recombinant LAMP-1 ectodomain in pH 5.0 running buffer, after which the sensor was dipped in pH 5.0 running buffer for 1200 s to measure LAMP-1 dissociation.

### Antibody quaternary preference assay using BLI

12.1F, 19.7E, 37.7H, and S370.7 IgGs were immobilized on AHC sensors (Sartorius) to a signal of 1.0 nM using an Octet Red96 instrument (ForteBio). The immobolized IgGs were then dipped in running buffer (PBS, 0.1% BSA, 0.02% Tween20, pH 7.4) followed by LIV GPC-I53-50A trimer, LIV GPC monomer, or running buffer. LIV GPC-I53-50A trimer and LIV GPC monomer were diluted in running buffer to concentrations that would contain the same amount of protomers in solution: 150 nM and 450 nM, respectively. Following a 600 s association period, the tips were dipped into running buffer and dissociation was measured for 600 s.

### Synthetic matriglycan microarray printing and screening

The synthesis of matriglycan compounds were reported previously (Sheikh et al., 2022). All compounds were printed on NHS-ester activated glass slides (NEXTERION® Slide H, Schott Inc.) using a Scienion sciFLEXARRAYER S3 non-contact microarray equipped with a Scienion PDC80 nozzle (Scienion Inc.). Individual compounds were dissolved in sodium phosphate buffer (0.225 M, pH 8.5) at the desired concentration and were printed in replicates of 6 with spot volume ∼ 400 pL, at 20°C and 50% humidity. Each slide has 24 subarrays in a 3×8 layout. After printing, slides were incubated in a humidity chamber for 8 hours and then blocked for one hour with a 5 mM ethanolamine in a Tris buffer (pH 9.0, 50 mM) at 50°C. Blocked slides were rinsed with DI water, spun dry, and kept in a desiccator at room temperature for future use.

Printed glass slide was pre-blocked with a solution of 1x TSM binding buffer (20 mM Tris·HCl, pH 7.4, 150 mM NaCl, 2 mM CaCl2, and 2 mM MgCl2, 0.05% Tween-20, 1% BSA) for 90 mins and the blocking solution was discarded. The Strep-tagged GPC-I53-50A containing the native site-1 protease cleavage site (1 μg/mL) was incubated with mAbs (5 μg/mL) in TSM binding buffer at 4°C for 1 h before StrepMAB-Classic Oyster 645 conjugate (0.5 μg/mL, IBA Lifesciences 2-1555-050) was added, and the solution was further incubated for another 30 min at 4°C. For the detection of the monoclonal antibody, a Cy3 conjugated goat-anti-human IgG antibody was used (5 μg/mL, Jackson Immuno Research, 109-165-008). The solution was then added to the microarray slide and the slide was incubated at room temperature for 1 h. The slide was sequentially washed with TSM wash buffer (20 mM Tris·HCl, pH 7.4, 150 mM NaCl, 2 mM CaCl2, and 2 mM MgCl2, 0.05% Tween-20), TSM buffer (20 mM Tris·HCl, pH 7.4, 150 mM NaCl, 2 mM CaCl2, and 2 mM MgCl2) and water.

The slides were scanned using a GenePix 4000B microarray scanner (Molecular Devices) at the appropriate excitation wavelength with a resolution of 5 μM. Various gains and PMT values were employed in the scanning to ensure all the signals were within the linear range of the scanner’s detector and there was no saturation of signals. The image was analyzed using GenePix Pro 7 software (version 7.2.29.2, Molecular Devices). The data was analyzed with an Excel macro (https://doi.org/10.5281/zenodo.5146251) to provide the results. The highest and lowest value of the total fluorescence intensity of the six replicates spots were removed, and the four values in the middle were used to provide the mean value and standard deviation.

### Pseudovirus neutralization assay

LASV pseudoviruses were made as previously described (Brouwer et al., 2022; Robinson et al., 2016) and pseudovirus neutralization assays were also performed as previously described using LASV psuedotyped viruses and TZM-bI cells (Brouwer et al., 2022). IC_50_ values were determined as the concentration at which infectivity was inhibited by 50% using Prism 9 (GraphPad).

### GPC-Fab complex formation

Purified GPC-I53-50A was incubated with purified Fabs for at least 1 h at 4°C at a 1:9 molar ratio of GPC-I53-50A to Fab. Next, complexes were purified from unbound Fab by size exclusion chromatography using a Superdex 200 increase 10/300 GL column. Fractions corresponding to GPC-Fab complexes (9-10.5 mL) were pooled and concentrated using a MWCO concentrator with a cutoff of 100 kDa (Millipore).

### B-cell sorting

We used two GPC bait constructs for isolating LASV-specific B cells: LIV GPC-I53-50A and Josiah rGPe (Robinson et al. 2016) with a T4-foldon domain. Biotinylated antigens were barcoded by incubation with barcoding complexes (TotalSeq-C, BioLegend) at a 2:1 molar ratio, resulting in an average of 2 antigen molecules per antigen-barcode complex (AgBC). We separately produced two AgBCs for each antigen using different fluorophores (APC and PE) and different barcodes to allow more stringent FACS selection and downstream data analysis. Previously cryopreserved PBMCs from a Sierra Leonean survivor of Lassa Fever (donor 1102370) were first stained with a “dark” human serum albumin AgBC (containing a barcode oligo but no fluorophore) prior to labeling with barcoded antigen baits and a small panel of flow cytometry Abs (anti-CD19 and a dump channel containing anti-CD3 and anti-CD14). All B cells (CD19+CD3-CD14-) double-positive for APC and PE were bulk sorted using a FACSMelody cell sorter (Beckton Dickinson). Antigen-selected B cells were then immediately processed on a 10x Genomics Chromium Controller using Next GEM 5’ v2 reagents as previously described (Hurtado et al., 2022). The resulting single cell sequencing libraries (gene expression, feature barcode and VDJ-B) were sequenced on an Illumina NovaSeq 6000 using a 100-cycle SP v1.5 reagent kit. Raw sequencing data was processed with CellRanger (Zheng et al., 2017a) and antibody sequences were annotated using the ab[x] toolkit (Briney and Burton, 2018). Specificity classification was determined from AgBC data using scab (Hurtado et al., 2022).

## Acknowledgements

The authors thank Bill Anderson and Hannah Turner from The Scripps Research Institute for their help with electron microscopy experiments. We thank Lauren Holden and Gabriel Ozorowski for their help in preparing this paper. We also thank Thijn Brummelkamp and Lars Hangartner for kindly sharing the LAMP-1 and BirA plasmid, respectively. We kindly thank Robert F. Garry for help with sample acquisition and collection from the Lassa fever survivor cohort at the Kenema Government Hospital. H.R.P. is supported by a David C. Farichild Endowed Fellowship and the Achievement Rewards for College Scientists Foundation. P.J.M.B. is supported by a Rubicon fellowship from the Netherlands Organisation for Scientific Research (NWO). A.A. is supported by the amfAR Mathilde Krim Fellowship in Biomedical Research (#110182-69-RKVA). Further support from the Vici fellowship from the Netherlands Organisation for Scientific Research (NWO; to R.W.S.), by the Fondation Dormeur, Vaduz (to R.W.S.), NIH grant R01 AI165692 (to G.J.B.), the International AIDS Vaccine Initiative (IAVI) through grant INV-008352/OPP1153692 funded by the Bill and Melinda Gates Foundation (to M.C.), and the Bill and Melinda Gates Foundation through grant OPP1170236 (to A.B.W.) enabled this work.

## Author contributions

Conceptualization: HRP, PJMB, AA, ABW

Methodology: HRP, PJMB, JH, JAB, MLN, LL

Formal analysis: HRP, PJMB, JH, JAB, MLN, LL, JHB, AA

Investigation: HRP, PJMB, JH, JAB, MLN, LL, JHB, GG, TM

Resources: JSS, GJB, MC, RWS, BB, ABW

Data curation: HRP

Writing Original Draft: HRP, PJMB, ABW

Writing, Review & Edit: HRP, PJMB, JH, JAB, MLN, LL, JHB, GG, TM, JSS, AA, GJB, MC, RWS, BB, ABW

Visualization: HRP, JH, JAB, MLN, LL

Supervision: AA, GJB, MC, RWS, BB, ABW

Project admin: HRP, PJMB, JAB, GJB, MC, RWS, BB, ABW

Funding acquisition: GJB, MC, RWS, BB, ABW

## Declaration of interests

The authors declare no conflicting interests.

## Data availability

Maps generated from the electron microscopy data are deposited in the Electron Microscopy Databank (http://www.emdatabank.org/) under accession IDs EMD-28178, EMD-28170, EMD-28180, EMD-28181, EMD-28182, EMD-28183, and EMD-28184. Atomic models corresponding to these maps have been deposited in the Protein Data Bank (http://www.rcsb.org/) under accession IDs 8EJD, 8EJE, 8EJF, 8EJG, 8EJH, 8EJI, and 8EJJ. Mass spectrometry raw files have been deposited in the MassIVE proteomics619database.

## Notes

### Competing Interest Statement

The authors have declared no competing interest.

