## Supplemental Information for "Structural conservation of Lassa virus glycoproteins and recognition by neutralizing antibodies"

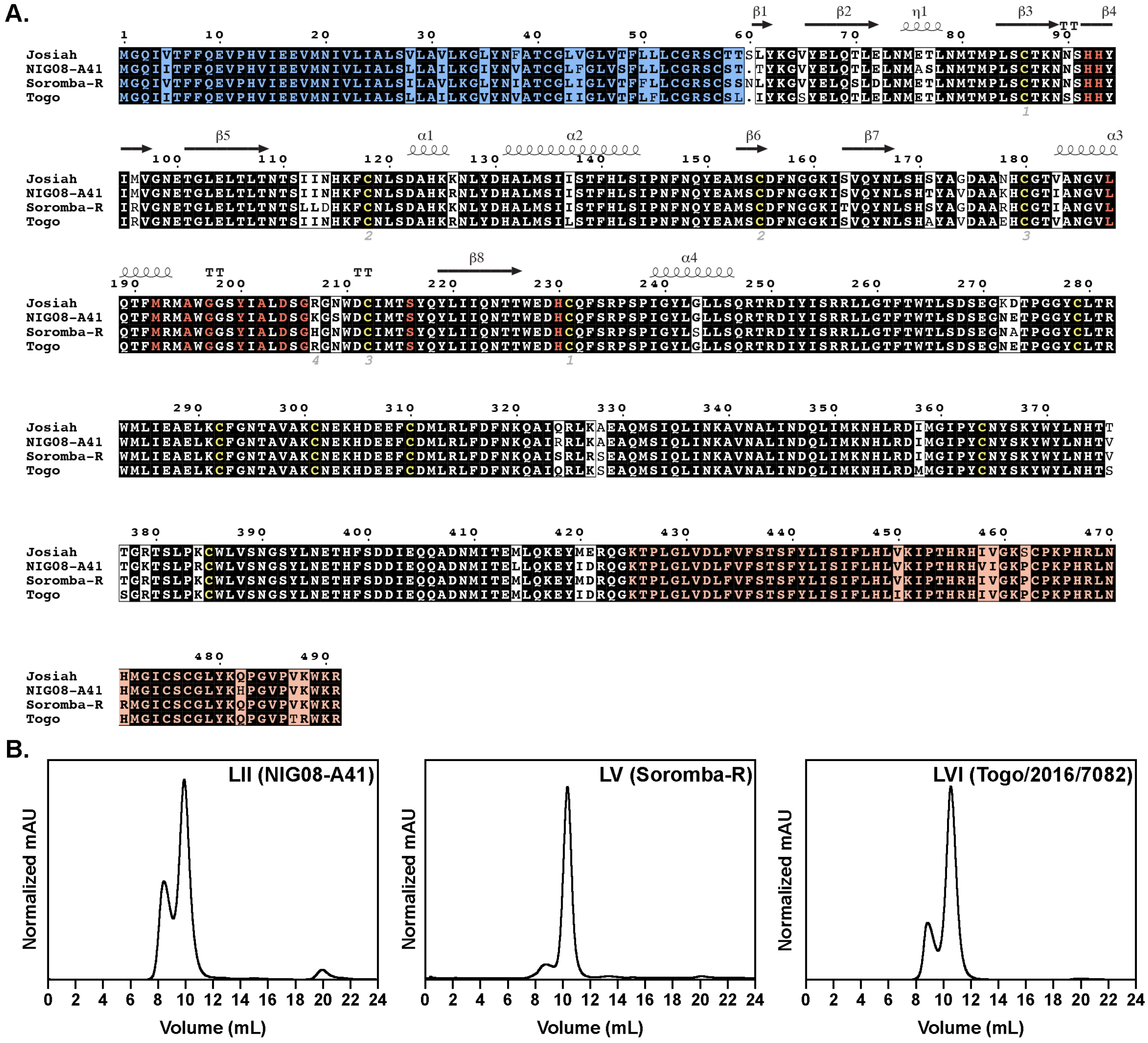


**Fig. S1: GPC lineage sequences and purification as trimers**. Related to Figure 1. (A) Native amino acid sequences from the four GPCs investigated in this study (lineage IV, Josiah; lineage II, NIG08-A41, lineage V, Soromba-R, and lineage VI, Togo) were aligned using the T-Coffee multiple sequence alignment server (Notredame et al., 2000). Residues shown as white text on a black background are conserved between the four strains. The stable signal peptide (SSP) is highlighted in blue and the transmembrane domain in orange. The histidine triad and additional residues required for LAMP-1 binding are highlighted in red (Cohen-Dvashi et al., 2015; Israeli et al., 2017). Ectodomain residues involved in disulfide bonds are highlighted yellow and numbered below. GPCysR4 stabilizing mutation (Hastie et al., 2017) locations are indicated with asterisks below. Secondary structural features are depicted above residues based on the atomic model for LIV GPC (PDB 8EJD). Accession codes for the sequences are as follows: Josiah, NP_694870.1; NIG08-A41, ADU56626.1; Soromba-R, AHC95553.1; and Togo/2016/7082, AMR44577.1. Sequence alignment visualized using ESPript3.0 (Robert and Gouet, 2014). (B) SEC chromatograms of LII, LV, and LVI GPC-I53-50As.

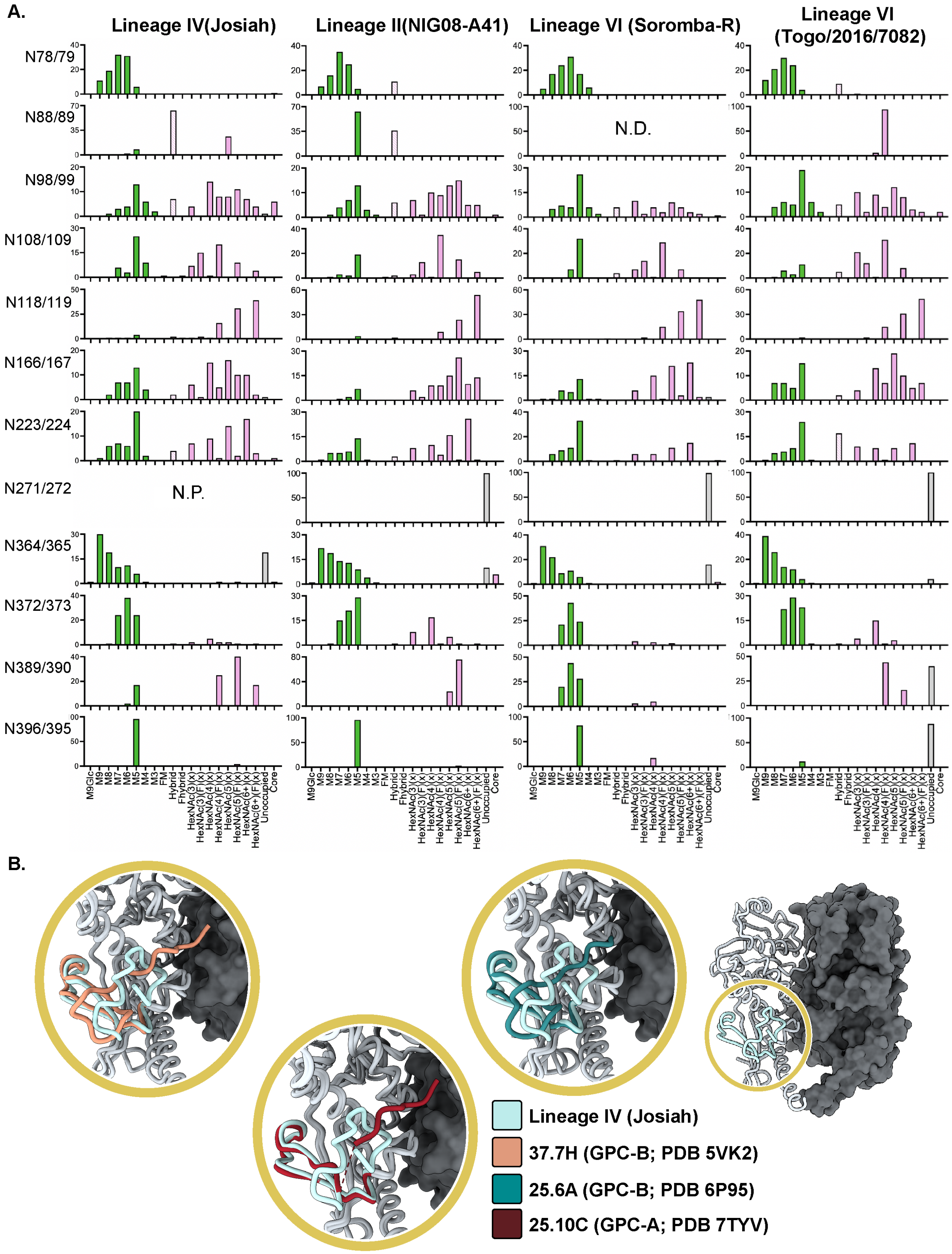


**Fig. S2: Detailed glycan and fusion peptide analyses.** Related to Figure 2. (A) Quantification and identities of glycan types determined by LCMs for each PNGS of each LASV lineage. Oligomannose-type glycans are shown in green, hybrid in dashed pink, complex glycans in pink and unoccupied sites in gray. N.D. indicates a PNGS which was undetected in the assay. N.P. indicates no PNGS is present in the sequence at that site. HexNAc(2)Hex(9−5) was classified as M9 to M3. Any of these structures containing a fucose were categorized as FM (fucosylated mannose). HexNAc(3)Hex(5−6)X was classified as Hybrid with HexNAc(3)Hex(5-6)Fuc(1)X classified as Fhybrid. Complex-type glycans were classified according to the number of HexNAc subunits and the presence or absence of fucosylation. As this fragmentation method does not provide linkage information, compositional isomers are grouped. For example, a triantennary glycan contains $HexNAc_5$ but so does a biantennary glycans with a bisect. Core glycans refer to truncated structures smaller than M3. M9Glc-M4 were classified as oligomannose-type glycans. (B) Ligand-free LIV GPC (PDB 8EJD) overlaid with additional GPC-B competition group abs 37.7H and 25.6A and GPC-A competition group ab 25.10C. Fusion peptides (residues 260-298) are colored for each model.

**A.**

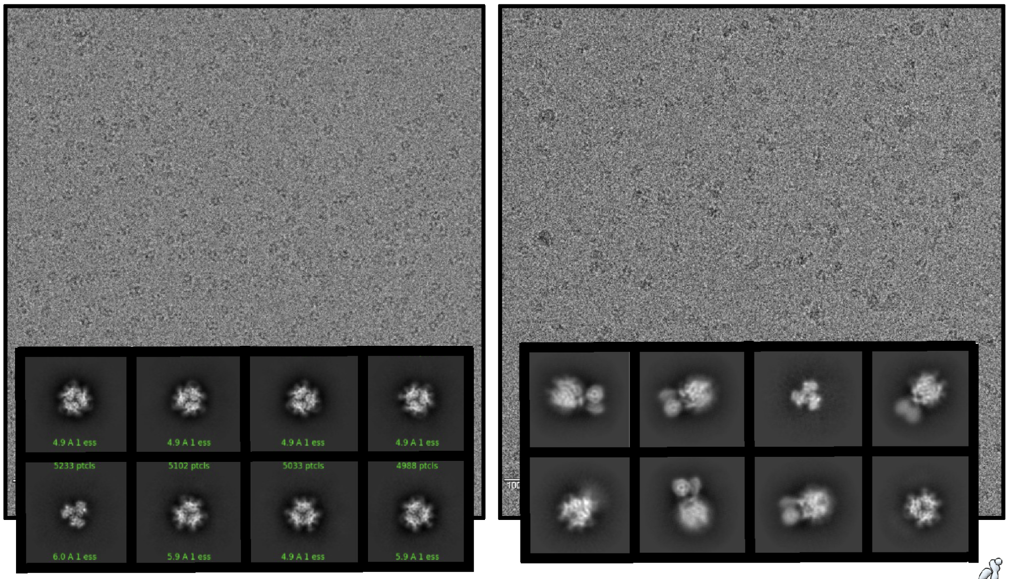

**B.**

**CryoSPARC**
1. Import micrographs
2. GCTF
3. Template picking & particle extraction
4. Iterative 2D classification
5. Heterogeneous refinement with scaffold - C1 symmetry
6. Homogeneous Refinment with scaffold- C1 symmetry
7. 3X Local Refinement masking out scaffold - C1 symmetry

**Relion**
8. Import particles
9. Local 3D refinement - C1 symmetry
10. 3D Classification - 3 classes, C1 symmetry
11. Local 3D refinement - C1 symmetry
12. Local 3D refinement - C3 symmetry
13. CTF Refinement
14. Local 3D refinement - C3 symmetry

**CryoSPARC**
15. Import particles
16. Local Refinement - C3 symmetry
17. Global CTF Refinement
18. Local refinement - C3 symmetry

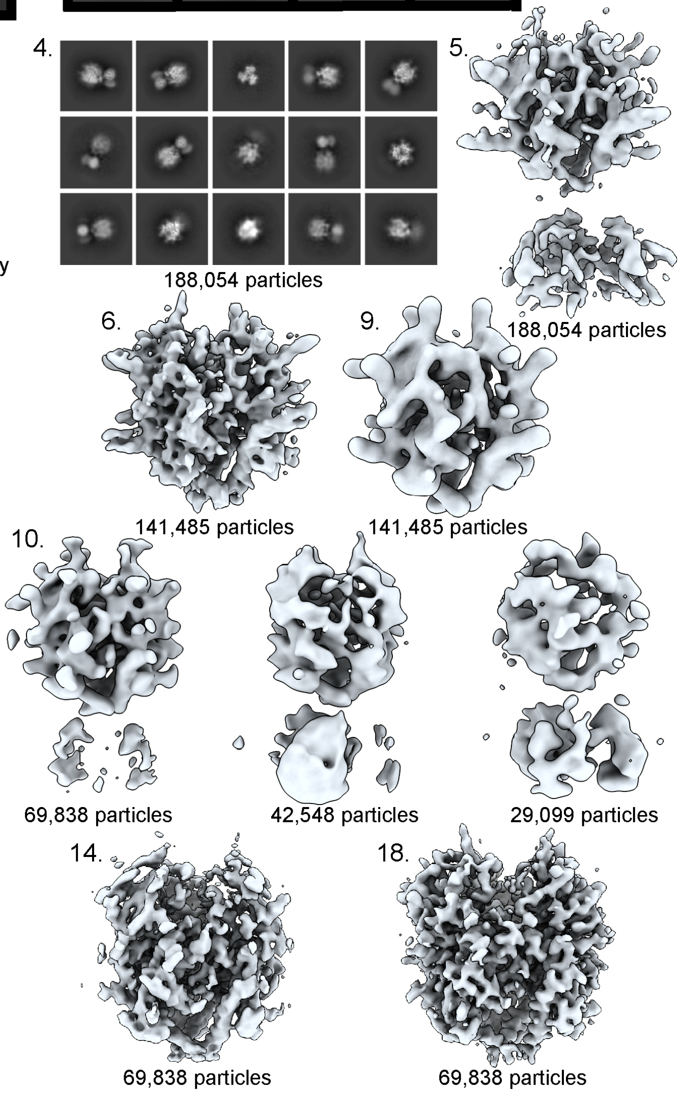


**Fig. S3: Optimizing single-particle cryoEM for GPC samples.** Related to Figure 2. (A) Sample micrographs showing effectiveness of fluoryl-octyl maltoside in improving GPC orientation in vitreous ice (right) compared to other detergents such as LMNG (left). (B) Representative data processing overview schematic showing data from EMD-28179. A similar processing scheme was employed for EMD- 28178, EMD-28180, and EMD-28181.

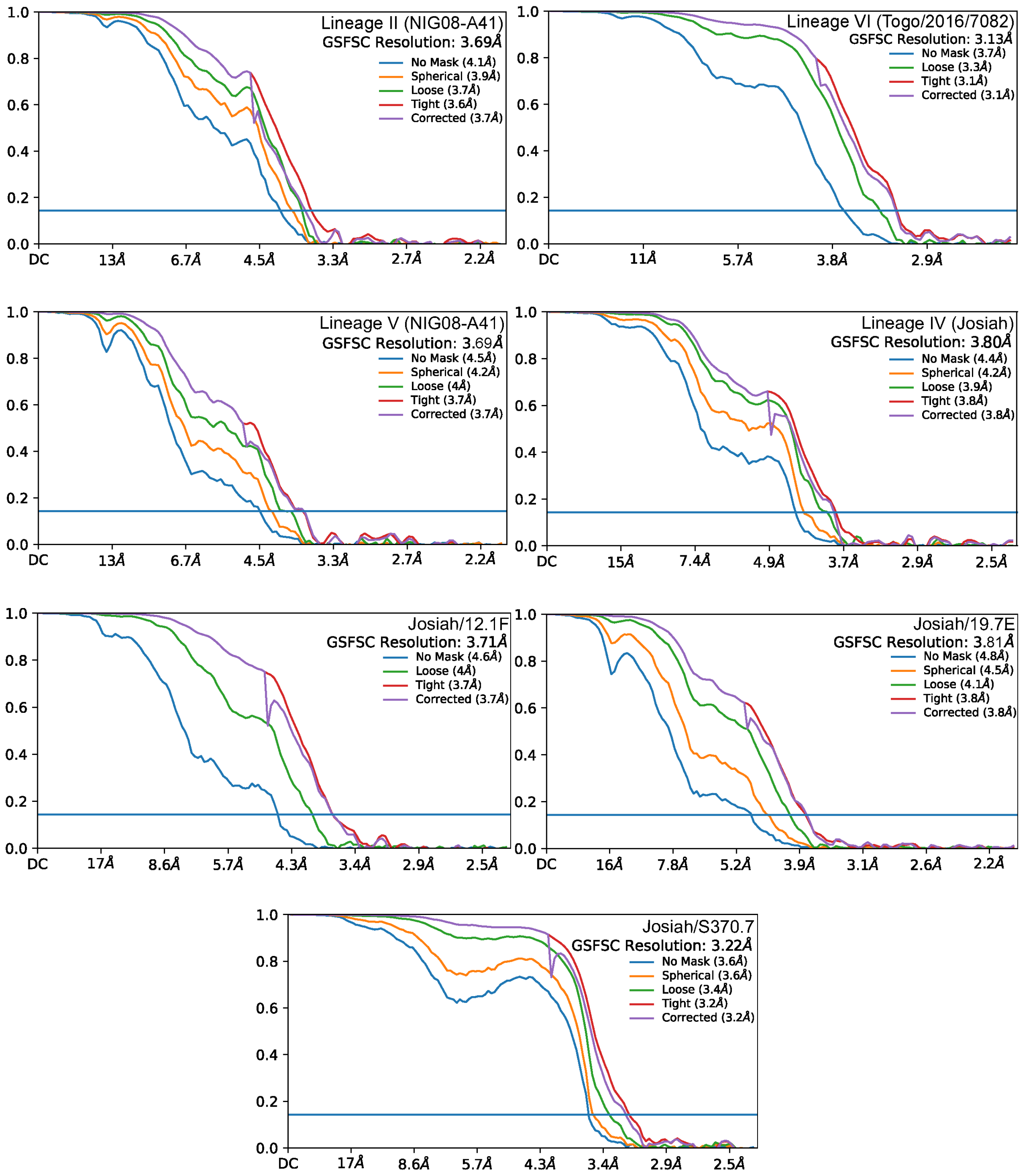


**Fig. S4: FSC plots for EM maps of LASV GPCs and GPCs bound to 12.1F, 19.7E, and S370.7.** Related to Figures 2, 4, and 6. Reported resolutions coincide with an FSC cutoff of 0.143. Plots were generated in cryoSPARC 3.2 (Punjani et al., 2017).

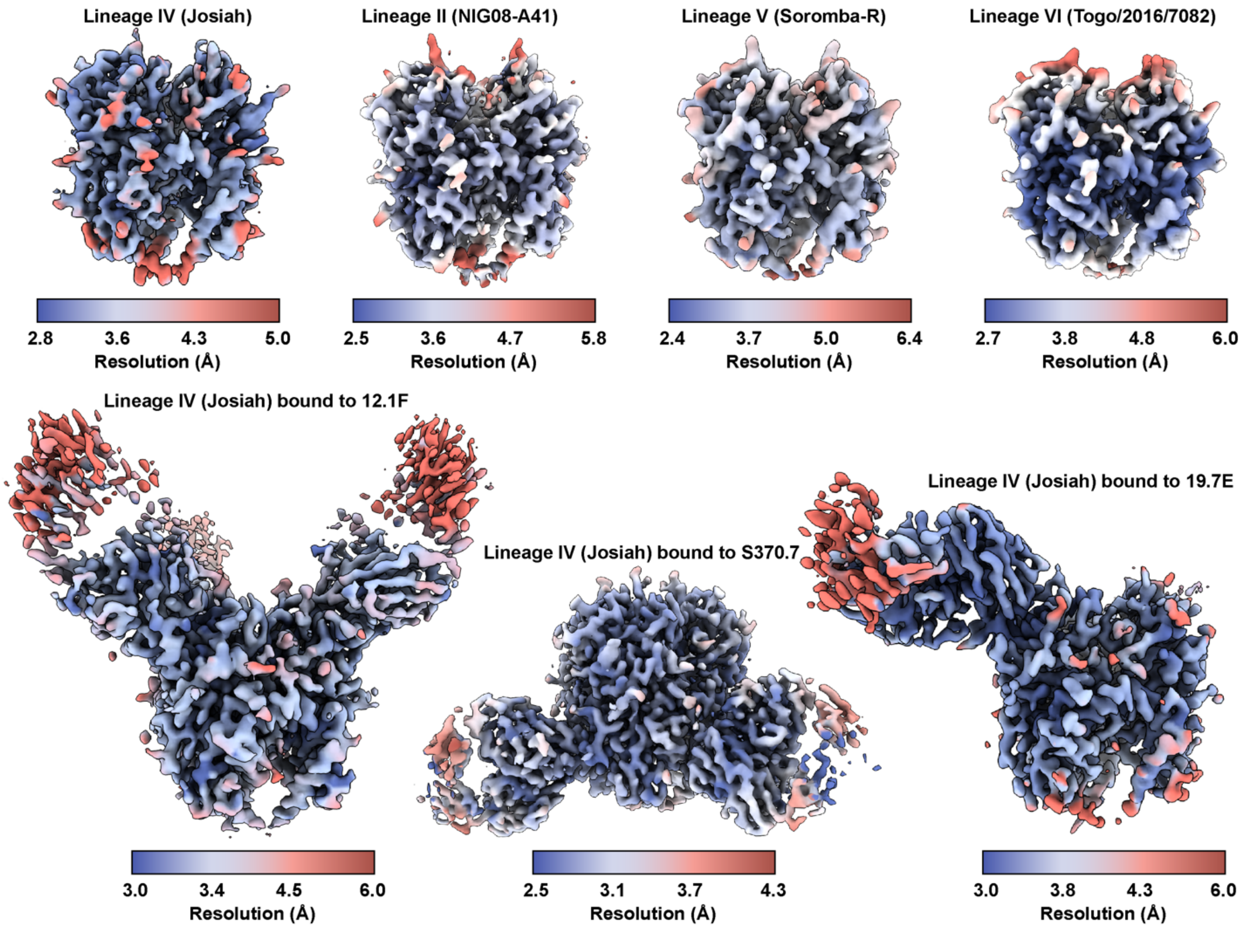


**Fig. S5: Local resolution plots for EM maps of LASV GPCs and GPCs bound to 12.1F, 19.7E, and S370.7.** Related to Figures 2, 4, and 6. Local resolution was calculated according to a 0.143 FSC threshold in cryoSPARC 3.2 (Punjani et al., 2017) and visualized in ChimeraX (Pettersen et al., 2021).

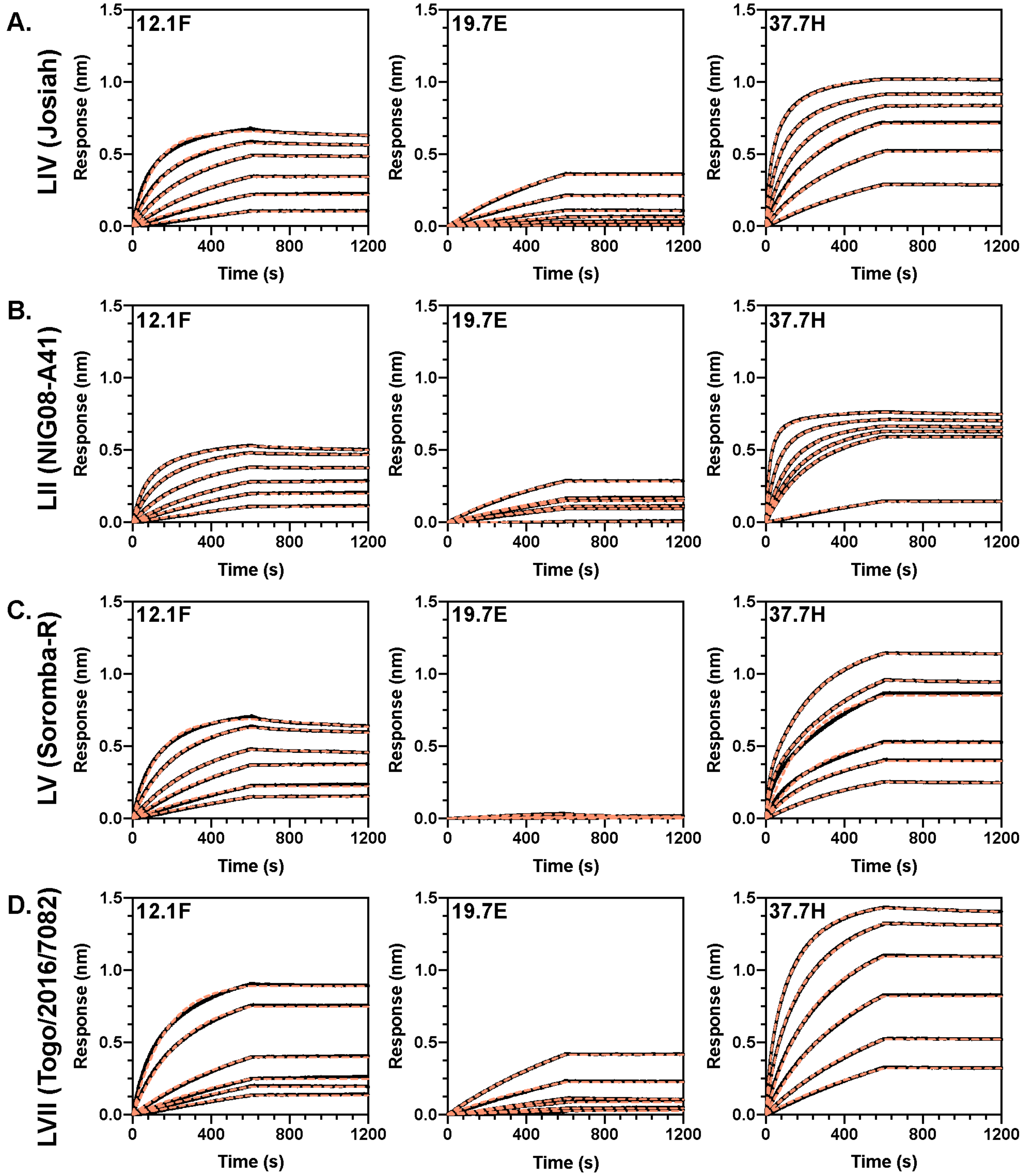


**Fig. S6: Binding profiles of GPC-I53-50A trimers to NAbs 12.1F, 19.7E, and 37.7H.** Related to Figure 3. BLI sensorgrams of immobilized biotinylated GPC-I53-50A binding to indicated IgGs at concentrations of 400, 200, 100, 50, 25, and 12.5 nM (black). Dotted orange lines represent the fit used to calculate on-rates and $R_{max}$ values.

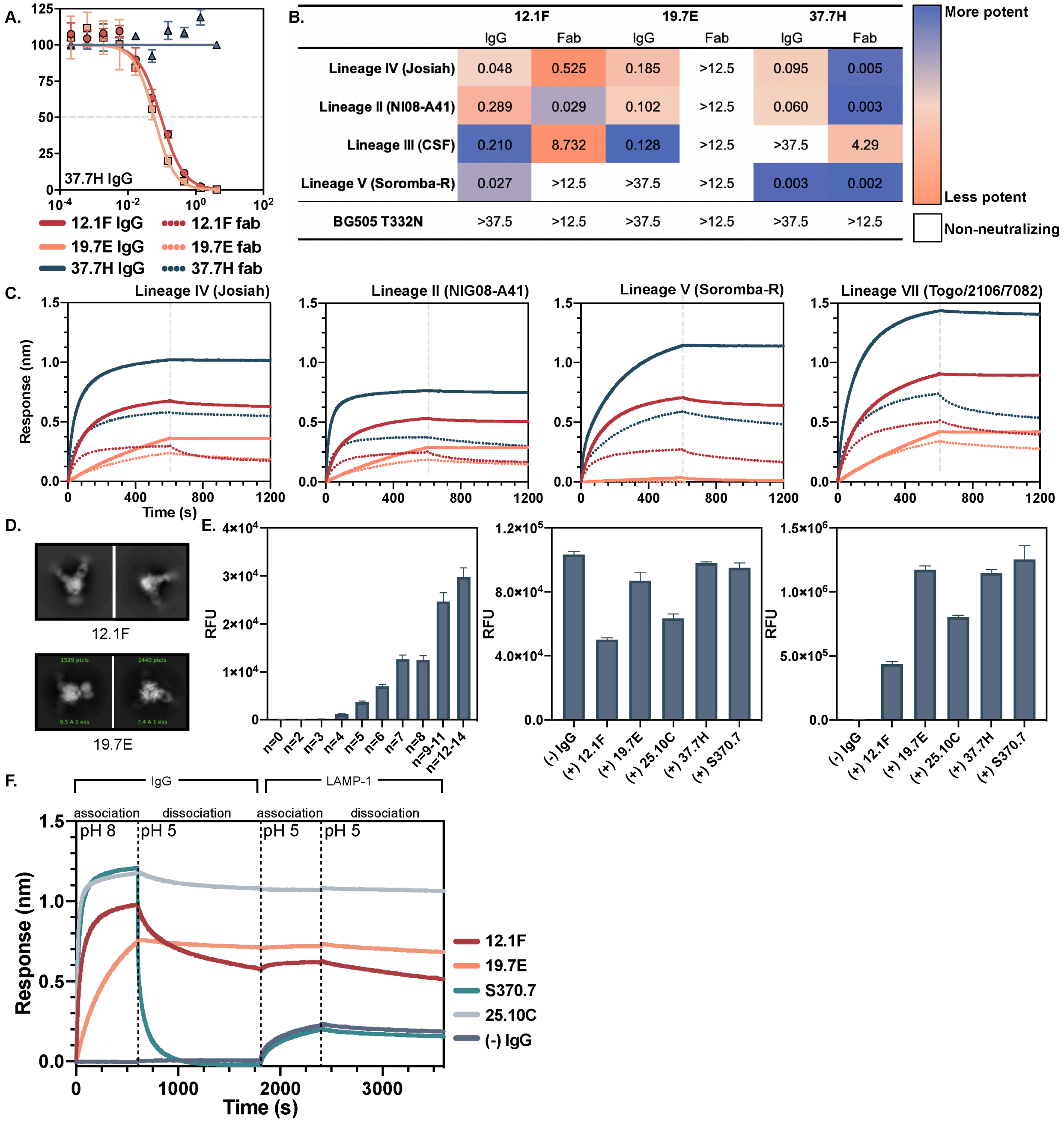


| | 12.1F | | 19.7E | | 37.7H | |
|---|---|---|---|---|---|---|
| | IgG | Fab | IgG | Fab | IgG | Fab |
| Lineage IV (Josiah) | 0.048 | 0.525 | 0.185 | >12.5 | 0.095 | 0.005 |
| Lineage II (NI08-A41) | 0.289 | 0.029 | 0.102 | >12.5 | 0.060 | 0.003 |
| Lineage III (CSF) | 0.210 | 8.732 | 0.128 | >12.5 | >37.5 | 4.29 |
| Lineage V (Soromba-R) | 0.027 | >12.5 | >37.5 | >12.5 | 0.003 | 0.002 |
| BG505 T332N | >37.5 | >12.5 | >37.5 | >12.5 | >37.5 | >12.5 |

**Fig. S7: 37.7H pseudovirus neutralization, IC$_{50}$ summary table, and comparison of binding between IgG and Fabs.** Related to Figure 3. (A) Pseudovirus neutralization of LASV lineages by 37.7H. The dotted line indicates 50% neutralization. Data points represent the mean with error bars indicating the SEM of three technical replicates. (B) IC$_{50}$ summary table of pseudovirus neutralization using 12.1F, 19.7E, and 37.7H IgG and Fabs. (C) BLI sensorgrams indicating binding behavior of immobilized GPC-I53-50A trimer binding to 400 nM of IgG or Fab. (D) Sample 2D classes from LIV GPCs binding to 12.1F (left) and 19.7E (right) showing preferred occupancies. While these occupancies were observed most often, it is important to note alternative occupancies were also observed in the data. (E) Matriglycan binding microarray controls. The StrepTagged GPC-I53-50A with a native site-1 protease cleavage site shows length-dependent binding to synthetic matriglycans on the microarray with *n* indicating the number of xylose and glucuronic acid disaccharide repeating units (left). The GPC-I53-50A trimers were detected by StrepMab antibody (middle). The monoclonal antibody used for detection was detected using anti-human Fc antibody (right). (F) Set-up for BLI LAMP-1 competition assay.

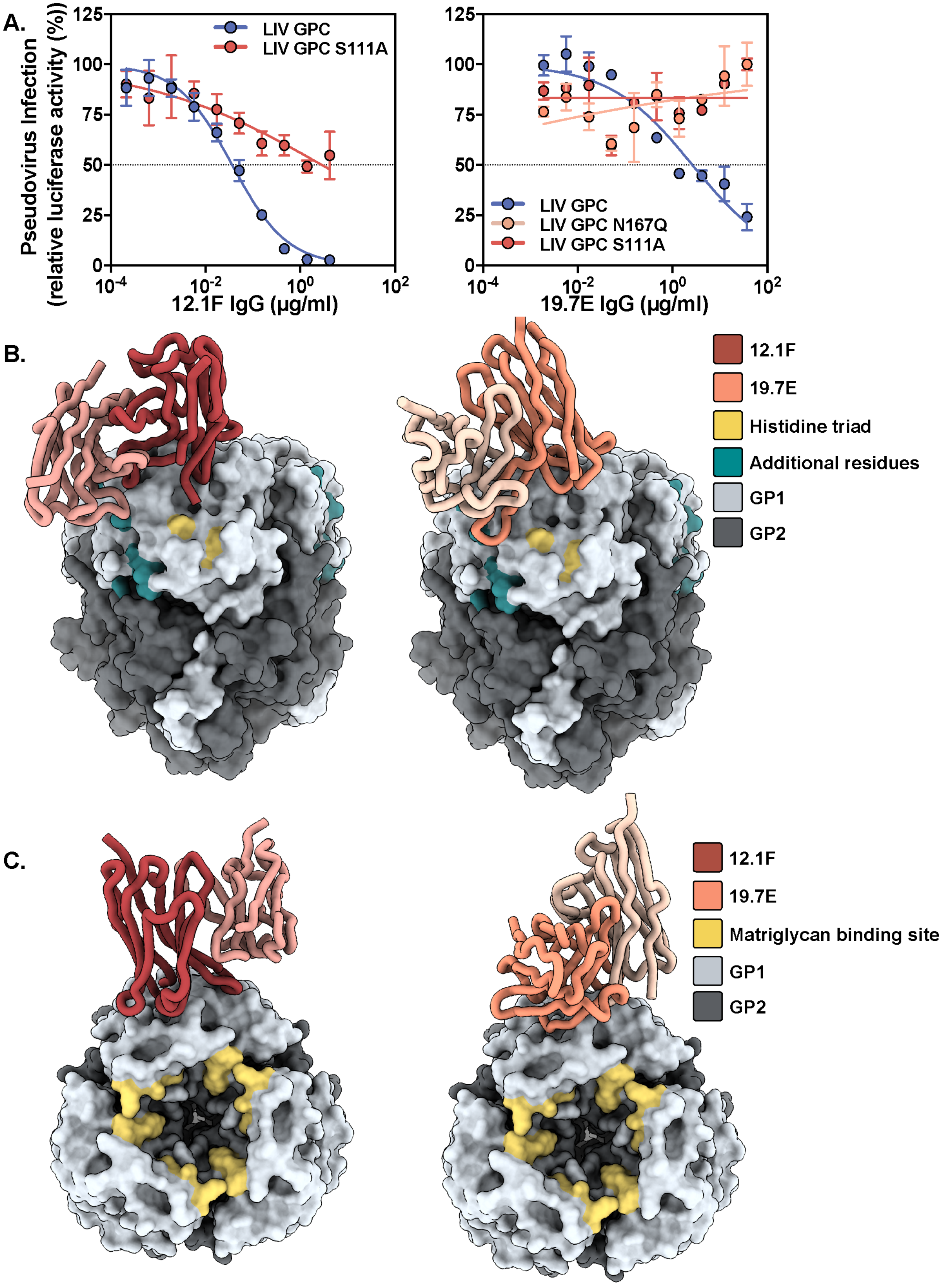


**Fig. S8: 12.1F and 19.7E glycan dependence and binding with respect to matriglycan and putative LAMP-1 binding sites.** Related to Figure 4. (A) Pseudovirus neutralization of LASV lineages by indicated mAbs to LIV pseudovirus containing the glycan knockout mutations S111A or N167Q. The dotted line indicates 50% neutralization. Data points represent the mean with error bars indicating the SEM of three (12.1F) or two (19.7E) technical replicates. (B) 12.1F and 19.7E Fabs overlaid on PDB 8EJD with residues important for LAMP-1 binding (Cohen-Dvashi et al., 2015; Israeli et al., 2017) shown in teal and gold. (B) 12.1F and 19.7E Fabs overlaid on PDB 8EJD with residues important for matriglycan binding shown in gold (Katz et al., 2022).

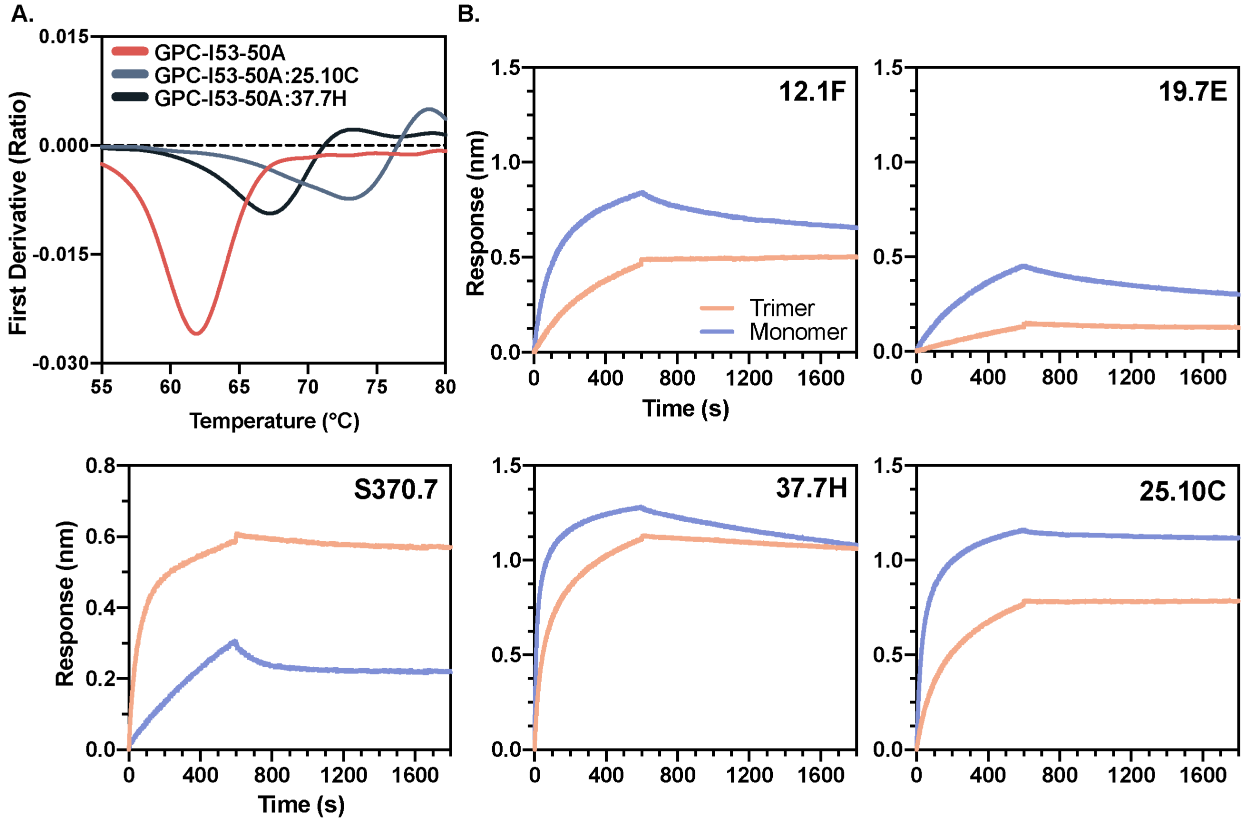


**Fig. S9: Biophysical comparison of S370.7 to known nAbs.** Related to Figure 6. (A) Thermostability of LASV Josiah-GPC in complex with S370.7 assessed by nanoDSF. Each melting curve is a representative of triplicate curves. (B) BLI sensorgrams comparing immobilized IgG binding to GPC-I53-50A trimer or GPC monomer. Trimer and monomer were diluted to 150 and 450 nM, respectively, which represents equivalent concentrations of protomer.

**Table S1: CryoEM map and atomic model refinement.** Related to Fig. 2, 4, and 6.

| | Lineage IV (Josiah) | Lineage II (NIG08-A41) | Lineage V (Soromba-R) | Lineage VI (Togo/2016/7082) | Josiah bound to 12.1F fab | Josiah bound to 19.7E fab | Josiah bound to S370.7 fab |
|---|---|---|---|---|---|---|---|
| **Access codes** | | | | | | | |
| PDB | 8EJD | 8EJE | 8EJF | 8EJG | 8EJH | 8EJI | 8EJJ |
| EMDB | EMD-28178 | EMD-28179 | EMD-28180 | EMD-28181 | EMD-28182 | EMD-28183 | EMD-28184 |
| Genebank | NP_694870.1 | ADU56626.1 | AHC95553.1 | AMR44577.1 | NP_694870.1 | NP_694870.1 | NP_694870.1 |
| **Data collection and processing** | | | | | | | |
| Microscope | Talos Arctica | Titan Krios | Titan Krios | Talos Arctica | Talos Arctica | Titan Krios | Talos Arctica |
| Magnification | 36,000 | 130,000 | 130,000 | 36,000 | 36,000 | 130,000 | 36000 |
| Voltage (kV) | 200 | 300 | 300 | 200 | 200 | 300 | 200 |
| Electron exposure ($e^-/Å^2$) | 50.0 | 49.2 | 49.2 | 50.3 | 50.3 | 50.2 | 50.1 |
| Defocus range (μm) | -0.7 to -2 | -0.7 to -2 | -0.7 to -2 | -0.7 to -2 | -0.7 to -2 | -0.7 to -2 | -0.7 to -2 |
| Pixel size (Å) | 1.150 | 1.045 | 1.045 | 1.150 | 1.150 | 1.045 | 1.150 |
| Imposed Symmetry | C3 | C3 | C3 | C3 | C3 | C1 | C3 |
| Final particle number | 56,496 | 69,838 | 27,663 | 71,504 | 62,262 | 70,071 (symmetry expanded) | 96,449 |
| Map resolution (Å) | 3.8 | 3.7 | 3.7 | 3.1 | 3.7 | 3.8 | 3.2 |
| FSC Threshold | 0.143 | 0.143 | 0.143 | 0.143 | 0.143 | 0.143 | 0.143 |
| Map sharpening B-factor ($Å^2$) | -70 | -93 | -67 | -50 | -130 | -120 | -63 |
| **Model refinement and validation** | | | | | | | |
| Total Residues | 1173 | 1134 | 1170 | 1185 | 1860 | 1335 | 1830 |
| Amino-acids | 1086 | 1068 | 1080 | 1083 | 1761 | 1257 | 1743 |
| Carbohydrates | 87 | 66 | 90 | 102 | 99 | 78 | 87 |
| RMSD Bonds | 0.021 | 0.022 | 0.023 | 0.024 | 0.020 | 0.021 | 0.023 |
| RMSD Angles | 1.78 | 2.11 | 1.95 | 1.87 | 1.75 | 1.81 | 1.77 |
| **Ramachandran** | | | | | | | |
| Outliers (%) | 0 | 0 | 0 | 0 | 0 | 0 | 0 |
| Allowed (%) | 4.5 | 6.0 | 9.0 | 6.4 | 5.7 | 3.9 | 2.8 |
| Favored (%) | 95.5 | 94.0 | 91.0 | 93.6 | 94.3 | 96.1 | 97.2 |
| Rotamer outliers (%) | 0 | 0 | 0 | 0 | 0 | 0 | 0 |
| Clash score | 2.04 | 2.39 | 1.89 | 0.88 | 2.50 | 3.31 | 1.73 |
| Molprobity score | 1.29 | 1.42 | 1.47 | 1.19 | 1.42 | 1.40 | 1.08 |
| FSC model (0/0.143/0.5) | 3.3/3.7/4.1 | 3.3/3.5/3.8 | 3.5/3.7/4.2 | 3.0/3.2/3.4 | 3.3/3.5/3.9 | 3.3/3.6/4.0 | 2.9/3.1/3.4 |
| EMRinger score | 2.51 | 2.46 | 2.04 | 3.32 | 2.12 | 2.44 | 3.74 |

**Table S2: 12.1F antibody interactions with GP1.** Related to Figure 4. Amino acid interactions at the 12.1F epitope-paratope were determined using the online-based Epitope Analyzer platform (Montiel-Garcia et al., 2022). Glycan contacts were assessed by finding close contacts (<4 Å) of the GPC glycans with 12.1F Fab using ChimeraX (Pettersen et al., 2021)

| GP Residue number | Amino acid | Atom | Fab Residue number | Amino Acid | Heavy or light chain | Distance (Å) | Interaction Type |
|---|---|---|---|---|---|---|---|
| 91 | SER | CB-CD2 | 111.1 | PHE | H | 3.4 | Van-der-Waals |
| 92 | HIS | CE1-CD2 | 111.1 | PHE | H | 3.9 | Van-der-Waals |
| 107 | LEU | O-CD2 | 111.1 | PHE | H | 3.8 | Van-der-Waals |
| 108 | THR | C-O | 111 | GLY | H | 3.5 | Van-der-Waals |
| 108 | THR | CG2-CE2 | 111.1 | PHE | H | 3.7 | Van-der-Waals |
| 109 | ASN | CA-O | 111 | GLY | H | 3.9 | Van-der-Waals |
| 109 | ASN | N-O | 111 | GLY | H | 2.9 | Hydro-Bond |
| 110 | THR | CG2-CA | 111 | GLY | H | 3.9 | Van-der-Waals |
| 110 | THR | CG2-CB | 109 | SER | H | 3.8 | Van-der-Waals |
| 111 | SER | O-CE2 | 38 | PHE | H | 3.9 | Van-der-Waals |
| 112 | ILE | C-ND2 | 57 | ASN | H | 3.7 | Van-der-Waals |
| 112 | ILE | C-OG | 64 | SER | H | 3.6 | Van-der-Waals |
| 112 | ILE | CG2-CD1 | 59 | LEU | H | 3.5 | Hydr-Phbc |
| 112 | ILE | O-ND2 | 57 | ASN | H | 2.8 | Hydro-Bond |
| 112 | ILE | O-OG | 64 | SER | H | 2.7 | Hydro-Bond |
| 113 | ILE | C-OG | 64 | SER | H | 3.4 | Van-der-Waals |
| 113 | ILE | N-OG | 64 | SER | H | 3.9 | Hydro-Bond |
| 114 | ASN | CA-OG | 64 | SER | H | 3.6 | Van-der-Waals |
| 114 | ASN | CB-O | 65 | THR | H | 3.6 | Van-der-Waals |
| 114 | ASN | N-OG | 64 | SER | H | 2.7 | Hydro-Bond |
| 114 | ASN | ND2-O | 65 | THR | H | 2.9 | Hydro-Bond |
| 156 | ASP | CG-CD1 | 59 | LEU | H | 3.6 | Van-der-Waals |
| 217 | TYR | CE2-N | 111.2 | ALA | H | 3.6 | Van-der-Waals |
| 217 | TYR | OH-C | 111.1 | PHE | H | 4.0 | Van-der-Waals |
| 217 | TYR | OH-N | 111.2 | ALA | H | 2.9 | Hydro-Bond |
| 110 | THR | C-OD1 | 109 | ASP | L | 4.0 | Van-der-Waals |
| 111 | SER | CB-NE1 | 114 | TRP | L | 4.0 | Van-der-Waals |
| 111 | SER | CB-OD1 | 109 | ASP | L | 3.6 | Van-der-Waals |
| 111 | SER | N-OD1 | 109 | ASP | L | 3.4 | Hydro-Bond |
| 111 | SER | OG-NE1 | 114 | TRP | L | 2.6 | Hydro-Bond |
| 111 | SER | OG-OD1 | 109 | ASP | L | 2.8 | Hydro-Bond |
| 111 | SER | OG-OD2 | 109 | ASP | L | 3.5 | Hydro-Bond |
| 112 | ILE | C-CZ2 | 114 | TRP | L | 3.8 | Van-der-Waals |
| 114 | ASN | CB-CD2 | 114 | TRP | L | 3.8 | Van-der-Waals |
| 114 | ASN | OD1-NE1 | 114 | TRP | L | 3.4 | Hydro-Bond |
| 219 | TYR | OH-CG | 109 | ASP | L | 3.6 | Van-der-Waals |
| 219 | TYR | OH-OD1 | 109 | ASP | L | 3.9 | Hydro-Bond |
| 219 | TYR | OH-OD2 | 109 | ASP | L | 3.0 | Hydro-Bond |
| N89 glycan, O3-C1 Man | | O6-CB | 4 | LEU | H | 3.7 | - |
| N89 glycan, O3-C1 Man | | O6-CG | 4 | LEU | H | 3.8 | - |
| N89 glycan, O3-C1 Man | | O6-CD1 | 4 | LEU | H | 3.9 | - |
| N89 glycan, O3-C1 Man | | O6-N | 4 | LEU | H | 3.7 | - |
| N89 glycan, O3-C1 Man | | O6-OE2 | 28 | GLU | H | 2.8 | - |
| N89 glycan, O3-C1 Man | | C6-OE2 | 28 | GLU | H | 3.5 | - |
| N89 glycan, O3-C1 Man | | O6-CB | 28 | GLU | H | 3.6 | - |
| N89 glycan, O3-C1 Man | | OG-CG | 28 | GLU | H | 3.8 | - |
| N89 glycan, O3-C1 Man | | C6-CG | 28 | GLU | H | 4.0 | - |
| N89 glycan, O3-C1 Man | | O5-OG | 29 | SER | H | 3.2 | - |
| N89 glycan, O3-C1 Man | | C6-CB | 29 | SER | H | 3.6 | - |
| N89 glycan, O3-C1 Man | | C6-OG | 29 | SER | H | 3.9 | - |
| N89 glycan, O3-C1 Man | | O5-CB | 29 | SER | H | 3.7 | - |
| N89 glycan, O4-C1 Man | | O3-CZ | 37 | PHE | H | 3.7 | - |
| N89 glycan, O4-C1 GlcNAc | | O4-OH | 108 | TYR | H | 2.9 | - |
| N89 glycan, O4-C1 GlcNAc | | O3-OH | 108 | TYR | H | 3.1 | - |
| N89 glycan, O4-C1 Man | | O2-CE1 | 108 | TYR | H | 3.2 | - |
| N89 glycan, O4-C1 Man | | O2-CZ | 108 | TYR | H | 3.3 | - |
| N89 glycan, O4-C1 GlcNAc | | O4-CZ | 108 | TYR | H | 3.4 | - |
| N89 glycan, O4-C1 Man | | O2-CD1 | 108 | TYR | H | 3.5 | - |
| N89 glycan, O4-C1 Man | | O2-CE2 | 108 | TYR | H | 3.7 | - |
| N89 glycan, O4-C1 Man | | O2-OH | 108 | TYR | H | 3.8 | - |
| N89 glycan, O4-C1 Man | | O2-CD2 | 108 | TYR | H | 4.0 | - |
| N89 glycan, O4-C1 GlcNAc | | C4-OH | 108 | TYR | H | 3.3 | - |
| N89 glycan, O4-C1 GlcNAc | | O4-CE2 | 108 | TYR | H | 3.7 | - |
| N89 glycan, O4-C1 GlcNAc | | C3-OH | 108 | TYR | H | 3.8 | - |
| N89 glycan, O4-C1 Man | | O2-CG2 | 108 | TYR | H | 3.9 | - |
| N89 glycan, O4-C1 GlcNAc | | O3-CZ | 108 | TYR | H | 3.9 | - |
| N89 glycan, ND2-C1 GlcNAc | | O3-OH | 110 | TYR | H | 2.9 | - |
| N89 glycan, ND2-C1 GlcNAc | | O3-CZ | 110 | TYR | H | 3.6 | - |
| N89 glycan, ND2-C1 GlcNAc | | O7-OH | 110 | TYR | H | 3.2 | - |
| N89 glycan, ND2-C1 GlcNAc | | O3-CE1 | 110 | TYR | H | 3.5 | - |
| N89 glycan, ND2-C1 GlcNAc | | C7-OH | 110 | TYR | H | 3.7 | - |
| N89 glycan, O4-C1 GlcNAc | | C6-CE1 | 110 | TYR | H | 3.7 | - |
| N89 glycan, ND2-C1 GlcNAc | | N2-OH | 110 | TYR | H | 3.9 | - |
| N109 glycan, ND2-C1 GlcNAc | | C8-O | 111.2 | ALA | H | 3.0 | - |
| N109 glycan, ND2-C1 GlcNAc | | O7-O | 111.2 | ALA | H | 2.9 | - |
| N109 glycan, ND2-C1 GlcNAc | | O7-O | 111.2 | ALA | H | 3.2 | - |
| N109 glycan, ND2-C1 GlcNAc | | C8-C | 111.2 | ALA | H | 3.9 | - |
| N109 glycan, ND2-C1 GlcNAc | | O3-CE3 | 112 | TRP | H | 3.9 | - |
| N109 glycan, O4-C1 GlcNAc | | C5-CE2 | 112 | TRP | H | 3.9 | - |
| N109 glycan, O4-C1 GlcNAc | | C6-NE1 | 112 | TRP | H | 4.0 | - |
| N109 glycan, ND2-C1 GlcNAc | | O7-ND2 | 112.1 | ASN | H | 3.9 | - |
| N109 glycan, ND2-C1 GlcNAc | | O7-CE3 | 112.3 | TRP | H | 3.8 | - |
| N109 glycan, O4-C1 GlcNAc | | C5-CZ2 | 112.3 | TRP | H | 3.9 | - |
| N89 glycan, O3-C1 Man | | O4-CG2 | 116 | ASP | H | 3.9 | - |
| N89 glycan, O3-C1 Man | | O3-CB | 116 | ASP | H | 3.6 | - |
| N89 glycan, O3-C1 Man | | O4-CB | 116 | ASP | H | 3.8 | - |
| N89 glycan, O3-C1 Man | | O4-CG2 | 117 | VAL | H | 3.2 | - |
| N89 glycan, O3-C1 Man | | O6-CG2 | 117 | VAL | H | 3.3 | - |
| N109 glycan, ND2-C1 GlcNAc | | O6-OG | 36 | SER | L | 3.9 | - |
| N109 glycan, ND2-C1 GlcNAc | | C6-OG | 36 | SER | L | 4.0 | - |
| N89 glycan, O3-C1 Man | | O2-OG1 | 69 | THR | L | 2.5 | - |
| N89 glycan, O3-C1 Man | | O2-CG2 | 69 | THR | L | 3.5 | - |
| N89 glycan, O3-C1 Man | | C2-CG2 | 69 | THR | L | 3.8 | - |
| N89 glycan, O6-C1 Man | | O2-CG2 | 69 | THR | L | 3.3 | - |
| N89 glycan, O3-C1 Man | | O2-CB | 69 | THR | L | 3.5 | - |
| N89 glycan, O3-C1 Man | | C2-OG1 | 69 | THR | L | 3.6 | - |
| N167 glycan, ND2-C1 GlcNAc | | C8-OD2 | 109 | ASP | L | 3.7 | - |

**Table S3: 19.7E antibody interactions with GP1.** Related to Figure 4. Amino acid interactions at the 19.7E epitope-paratope region were determined using the online-based Epitope Analyzer platform (Montiel-Garcia et al., 2022). Glycan contacts were assessed by finding close contacts (<4 Å) of the GPC glycans with 19.7E Fab using ChimeraX (Pettersen et al., 2021).

| GP Residue number | Amino acid | Atom | Fab Residue number | Amino Acid | Heavy or light chain | Distance (Å) | Interaction Type |
|---|---|---|---|---|---|---|---|
| 92 | HIS | CE1-OE2 | 1 | GLU | H | 3.5 | Van-der-Waals |
| 104 | GLU | CD-OE1 | 1 | GLU | H | 3.8 | Van-der-Waals |
| 109 | ASN | C-CB | 114 | SER | H | 3.7 | Van-der-Waals |
| 109 | ASN | C-OD1 | 112 | ASP | H | 3.8 | Van-der-Waals |
| 109 | ASN | CB-CB | 113 | TRP | H | 4.0 | Van-der-Waals |
| 109 | ASN | ND2-O | 113 | TRP | H | 3.8 | Hydro-Bond |
| 109 | ASN | O-N | 113 | TRP | H | 3.4 | Hydro-Bond |
| 109 | ASN | O-N | 114 | SER | H | 2.9 | Hydro-Bond |
| 110 | THR | C-OD2 | 112 | ASP | H | 3.7 | Van-der-Waals |
| 110 | THR | CG2-CB | 114 | SER | H | 3.7 | Van-der-Waals |
| 111 | SER | C-OH | 37 | TYR | H | 3.5 | Van-der-Waals |
| 111 | SER | CA-OD2 | 112 | ASP | H | 3.6 | Van-der-Waals |
| 111 | SER | N-OD1 | 112 | ASP | H | 3.8 | Hydro-Bond |
| 111 | SER | N-OD2 | 112 | ASP | H | 2.8 | Hydro-Bond |
| 111 | SER | O-CZ | 107 | ARG | H | 3.7 | Van-der-Waals |
| 111 | SER | O-NH1 | 107 | ARG | H | 3.5 | Hydro-Bond |
| 111 | SER | O-NH2 | 107 | ARG | H | 3.1 | Hydro-Bond |
| 111 | SER | O-OH | 37 | TYR | H | 3.7 | Hydro-Bond |
| 111 | SER | OG-NH1 | 107 | ARG | H | 3.8 | Hydro-Bond |
| 111 | SER | OG-OD2 | 112 | ASP | H | 2.8 | Hydro-Bond |
| 111 | SER | OG-OH | 37 | TYR | H | 2.8 | Hydro-Bond |
| 112 | ILE | C-CE1 | 37 | TYR | H | 3.7 | Van-der-Waals |
| 112 | ILE | CA-CB | 28 | PHE | H | 3.9 | Hydr-Phbc |
| 112 | ILE | CB-CB | 28 | PHE | H | 3.9 | Hydr-Phbc |
| 112 | ILE | CG2-CG2 | 2 | VAL | H | 3.9 | Hydr-Phbc |
| 112 | ILE | CG2-O | 27 | GLY | H | 3.9 | Van-der-Waals |
| 112 | ILE | N-OH | 37 | TYR | H | 3.4 | Hydro-Bond |
| 112 | ILE | O-CA | 29 | SER | H | 3.7 | Van-der-Waals |
| 112 | ILE | O-N | 29 | SER | H | 2.9 | Hydro-Bond |
| 112 | ILE | O-OG | 29 | SER | H | 3.8 | Hydro-Bond |
| 112 | ILE | O-OH | 37 | TYR | H | 3.4 | Hydro-Bond |
| 113 | ILE | C-OH | 37 | TYR | H | 3.4 | Van-der-Waals |
| 113 | ILE | CG1-OG | 29 | SER | H | 3.5 | Van-der-Waals |
| 113 | ILE | N-OH | 37 | TYR | H | 3.0 | Hydro-Bond |
| 113 | ILE | O-OH | 37 | TYR | H | 3.8 | Hydro-Bond |
| 114 | ASN | CB-CE2 | 37 | TYR | H | 3.7 | Van-der-Waals |
| 114 | ASN | CG-C | 36 | SER | H | 4.0 | Van-der-Waals |
| 114 | ASN | N-OH | 37 | TYR | H | 3.4 | Hydro-Bond |
| 114 | ASN | ND2-O | 36 | SER | H | 3.0 | Hydro-Bond |
| 114 | ASN | OD1-N | 37 | TYR | H | 3.9 | Hydro-Bond |
| 114 | ASN | OD1-OH | 37 | TYR | H | 4.0 | Hydro-Bond |
| 115 | HIS | CB-OG | 36 | SER | H | 3.8 | Van-der-Waals |
| 115 | HIS | ND1-OG | 36 | SER | H | 3.7 | Hydro-Bond |
| 156 | ASP | CB-OG | 29 | SER | H | 3.4 | Van-der-Waals |
| 156 | ASP | OD2-OG | 29 | SER | H | 2.8 | Hydro-Bond |
| 169 | SER | CB-CZ2 | 113 | TRP | H | 4.0 | Van-der-Waals |
| 172 | TYR | CE1-CH2 | 113 | TRP | H | 4.0 | Hydr-Phbc |
| 216 | SER | O-CZ3 | 113 | TRP | H | 4.0 | Van-der-Waals |
| 217 | TYR | C-CE3 | 113 | TRP | H | 4.0 | Van-der-Waals |
| 218 | GLN | CA-CE3 | 113 | TRP | H | 3.8 | Van-der-Waals |
| 218 | GLN | NE2-O | 111 | TYR | H | 3.1 | Hydro-Bond |
| 218 | GLN | OE1-C | 112 | ASP | H | 3.7 | Van-der-Waals |
| 218 | GLN | OE1-N | 113 | TRP | H | 2.8 | Hydro-Bond |
| 223 | GLN | NE2-OE1 | 1 | GLU | H | 3.3 | Hydro-Bond |
| 223 | GLN | OE1-O | 27 | GLY | H | 3.5 | Van-der-Waals |
| N89 glycan, O4-C1 GlcNAc | | O7-CD | 1 | GLU | H | 4.0 | - |
| N89 glycan, O4-C1 GlcNAc | | O7-CG | 1 | GLU | H | 3.2 | - |
| N167, O4-C1 GlcNac | | O6-CE1 | 111 | TYR | H | 3.3 | - |
| N167, O4-C1 GlcNac | | O6-CD1 | 111 | TYR | H | 3.5 | - |
| N167, O4-C1 GlcNac | | O7-CD2 | 111 | TYR | H | 3.6 | - |
| N167 glycan, O4-C1 Man | | C1-OH | 111 | TYR | H | 3.7 | - |
| N167 glycan, O4-C1 Man | | C3-OH | 111 | TYR | H | 3.7 | - |
| N167, O4-C1 GlcNac | | C2-CD2 | 111 | TYR | H | 3.9 | - |
| N109 glycan, ND2-C1 GlcNAc | | C1-CE3 | 113 | TRP | H | 3.7 | - |
| N109 glycan, ND2-C1 GlcNAc | | O6-CD2 | 113 | TRP | H | 3.8 | - |
| N109 glycan, ND2-C1 GlcNAc | | O6-CE2 | 113 | TRP | H | 3.8 | - |
| N109 glycan, ND2-C1 GlcNAc | | O6-CD1 | 113 | TRP | H | 3.9 | - |
| N109 glycan, ND2-C1 GlcNAc | | O5-CE3 | 113 | TRP | H | 3.9 | - |
| N109 glycan, ND2-C1 GlcNAc | | O6-NE1 | 113 | TRP | H | 3.9 | - |
| N109 glycan, ND2-C1 GlcNAc | | O5-CB | 113 | TRP | H | 3.8 | - |
| N109 glycan, ND2-C1 GlcNAc | | O6-CG | 113 | TRP | H | 3.8 | - |
| N109 glycan, ND2-C1 GlcNAc | | C1-CB | 113 | TRP | H | 4.0 | - |
| N89 glycan, O6-C1 Man | | O4-CG | 118 | TRP | H | 3.6 | - |
| N89 glycan, O6-C1 Man | | O4-CD1 | 118 | TRP | H | 3.9 | - |
| N89 glycan, O6-C1 Man | | O4-CB | 118 | TRP | H | 3.6 | - |
| N89 glycan, O6-C1 Man | | C6-CB | 49 | ALA | L | 3.7 | - |
| N89 glycan, O6-C1 Man | | O6-CB | 49 | ALA | L | 3.8 | - |
| N89 glycan, O6-C1 Man | | O2-CG | 51 | LYS | L | 3.8 | - |
| N167 glycan, O3-C1 Man | | C6-OD1 | 108 | ASN | L | 3.1 | - |
| N167 glycan, O3-C1 Man | | O4-C | 108 | ASN | L | 3.5 | - |
| N167 glycan, O3-C1 Man | | C4-O | 108 | ASN | L | 3.8 | - |
| N167 glycan, O3-C1 Man | | O6-OD1 | 108 | ASN | L | 3.9 | - |
| N167 glycan, O3-C1 Man | | C6-CG | 108 | ASN | L | 4.0 | - |

**Table S4: S370.7 antibody interactions with GPC.** Related to Figure 4. Amino acid interactions at the S370.7 epitope-paratope region were determined using the online-based Epitope Analyzer platform (Montiel-Garcia et al., 2022). Glycan contacts were assessed by finding close contacts (<4 A) of the GPC glycans with 19.7E Fab using ChimeraX (Pettersen et al., 2021).

| GP Residue number | Amino acid | Atom | Fab Residue number | Amino Acid | Heavy or light chain | Distance (Å) | Interaction Type |
|---|---|---|---|---|---|---|---|
| 268 | ASP | CB-CG | 112.1 | PRO | H | 4.0 | Van-der-Waals |
| 268 | ASP | OD2-CB | 111.1 | SER | H | 3.9 | Van-der-Waals |
| 325 | ARG | NH1-CB | 112.4 | VAL | H | 3.7 | Van-der-Waals |
| 325 | ARG | NH1-O | 112.4 | VAL | H | 3.7 | Hydro-Bond |
| 325 | ARG | NH1-OG | 64 | SER | H | 3.8 | Hydro-Bond |
| 325 | ARG | NH2-O | 112.4 | VAL | H | 3.8 | Hydro-Bond |
| 360 | CYS | O-CG1 | 112.4 | VAL | H | 3.6 | Van-der-Waals |
| 362 | PRO | CD-CD2 | 112.5 | TYR | H | 3.9 | Hydr-Phbc |
| 362 | PRO | CG-CD2 | 112.5 | TYR | H | 3.4 | Hydr-Phbc |
| 362 | PRO | CG-CE2 | 112.5 | TYR | H | 3.8 | Hydr-Phbc |
| 387 | LEU | CD1-CG1 | 111.2 | VAL | H | 3.7 | Hydr-Phbc |
| 387 | LEU | CD1-CG2 | 111.2 | VAL | H | 4.0 | Hydr-Phbc |
| 389 | SER | OG-CE2 | 112.5 | TYR | H | 3.9 | Van-der-Waals |
| 394 | LEU | CD2-CE1 | 112.5 | TYR | H | 3.9 | Hydr-Phbc |
| 394 | LEU | CD2-CE2 | 112.5 | TYR | H | 3.9 | Hydr-Phbc |
| 394 | LEU | CD2-CZ | 112.5 | TYR | H | 3.6 | Hydr-Phbc |
| 397 | THR | O-CG | 111 | ARG | H | 3.8 | Van-der-Waals |
| 398 | HIS | C-C | 111 | ARG | H | 3.8 | Van-der-Waals |
| 398 | HIS | CD2-CD2 | 58 | HIS | H | 3.9 | Van-der-Waals |
| 398 | HIS | CG-CE1 | 112.5 | TYR | H | 3.9 | Van-der-Waals |
| 398 | HIS | O-C | 111.1 | SER | H | 3.4 | Van-der-Waals |
| 398 | HIS | O-N | 111.1 | SER | H | 3.5 | Hydro-Bond |
| 399 | PHE | C-O | 111 | ARG | H | 3.1 | Van-der-Waals |
| 399 | PHE | CB-CG2 | 111.2 | VAL | H | 3.8 | Hydr-Phbc |
| 399 | PHE | N-O | 111 | ARG | H | 3.5 | Hydro-Bond |
| 400 | SER | C-O | 111 | ARG | H | 3.8 | Van-der-Waals |
| 400 | SER | N-O | 111 | ARG | H | 3.0 | Hydro-Bond |
| 401 | ASP | CA-O | 111 | ARG | H | 3.6 | Van-der-Waals |
| 401 | ASP | N-O | 111 | ARG | H | 2.9 | Hydro-Bond |
| 401 | ASP | OD2-N | 111 | ARG | H | 3.7 | Hydro-Bond |
| 402 | ASP | CG-CB | 111.1 | SER | H | 4.0 | Van-der-Waals |
| 402 | ASP | CG-N | 111.2 | VAL | H | 3.9 | Van-der-Waals |
| 402 | ASP | OD2-N | 111.2 | VAL | H | 2.9 | Hydro-Bond |
| 269 | SER | OG-CE | 36 | LYS | L | 3.3 | Van-der-Waals |
| 269 | SER | OG-NZ | 36 | LYS | L | 3.0 | Hydro-Bond |
| 270 | GLU | O-CG | 36 | LYS | L | 3.4 | Van-der-Waals |
| 272 | LYS | CA-OH | 38 | TYR | L | 3.9 | Van-der-Waals |
| 272 | LYS | CB-O | 29 | PRO | L | 4.0 | Van-der-Waals |
| 272 | LYS | CD-C | 37 | GLN | L | 3.8 | Van-der-Waals |
| 272 | LYS | CD-O | 28 | LEU | L | 4.0 | Van-der-Waals |
| 272 | LYS | CD-OD2 | 57 | ASP | L | 3.4 | Van-der-Waals |
| 272 | LYS | CG-C | 36 | LYS | L | 3.9 | Van-der-Waals |
| 272 | LYS | N-O | 29 | PRO | L | 3.5 | Hydro-Bond |
| 272 | LYS | N-O | 36 | LYS | L | 3.9 | Hydro-Bond |
| 272 | LYS | NZ-O | 28 | LEU | L | 2.6 | Hydro-Bond |
| 272 | LYS | NZ-O | 37 | GLN | L | 3.1 | Hydro-Bond |
| 272 | LYS | NZ-OD1 | 57 | ASP | L | 3.6 | Salt-Bridge |
| 272 | LYS | NZ-OD2 | 57 | ASP | L | 2.8 | Salt-Bridge |
| 272 | LYS | O-OH | 38 | TYR | L | 3.5 | Hydro-Bond |
| 279 | CYS | CB-CD | 36 | LYS | L | 3.9 | Van-der-Waals |
| 281 | THR | CB-OD1 | 110 | ASP | L | 3.5 | Van-der-Waals |
| 281 | THR | CG2-C | 109 | SER | L | 3.9 | Van-der-Waals |
| 281 | THR | CG2-NZ | 36 | LYS | L | 3.8 | Van-der-Waals |
| 281 | THR | OG1-OD1 | 110 | ASP | L | 2.9 | Hydro-Bond |
| 283 | TRP | CH2-CB | 109 | SER | L | 3.9 | Van-der-Waals |
| 283 | TRP | CZ3-CG | 110 | ASP | L | 3.9 | Van-der-Waals |
| 290 | LEU | CD2-CB | 29 | PRO | L | 4.0 | Hydr-Phbc |
| 290 | LEU | CD2-CB | 36 | LYS | L | 3.9 | Van-der-Waals |
| 290 | LEU | CD2-O | 26 | ASP | L | 3.9 | Van-der-Waals |
| 320 | LYS | CE-O | 110 | ASP | L | 4.0 | Van-der-Waals |
| 320 | LYS | NZ-O | 110 | ASP | L | 3.8 | Hydro-Bond |
| 324 | GLN | NE2-OG1 | 114 | THR | L | 3.1 | Hydro-Bond |
| N390 glycan, ND2-C1 GlcNAc | | C8-O | 59 | SER | H | 3.2 | - |
| N390 glycan, ND2-C1 GlcNAc | | C8-C | 59 | SER | H | 3.8 | - |
| N390 glycan, ND2-C1 GlcNAc | | C8-CA | 63 | GLY | H | 4.1 | - |
| N79 glycan, O3-C1 Man | | O4-CE2 | 115 | TYR | L | 3.2 | - |
| N79 glycan, O3-C1 Man | | O4-CD2 | 115 | TYR | L | 3.4 | - |
| N79 glycan, O3-C1 Man | | C5-N | 2 | TYR | L | 3.9 | - |
| N79 glycan, O3-C1 Man | | O4-N | 2 | TYR | L | 3.0 | - |
